# Testing for phylogenetic signal in single-cell RNA-seq data

**DOI:** 10.1101/2021.01.07.425804

**Authors:** Jiří C. Moravec, Rob Lanfear, David L. Spector, Sarah D. Diermeier, Alex Gavryushkin

## Abstract

Phylogenetic methods are emerging as a useful tool to understand cancer evolutionary dynamics, including tumor structure, heterogeneity, and progression. Most currently used approaches utilize either bulk whole genome sequencing (WGS) or single-cell DNA sequencing (scDNA-seq) and are based on calling copy number alterations and single nucleotide variants (SNVs). scRNA-seq is commonly applied to explore differential gene expression of cancer cells throughout tumor progression. The method exacerbates the single-cell sequencing problem of low yield per cell with uneven expression levels. This accounts for low and uneven sequencing coverage and makes SNV detection and phylogenetic analysis challenging. In this paper, we demonstrate for the first time that scRNA-seq data contains sufficient evolutionary signal and can also be utilized in phylogenetic analyses. We explore and compare results of such analyses based on both expression levels and SNVs called from scRNA-seq data. Both techniques are shown to be useful for reconstructing phylogenetic relationships between cells, reflecting the clonal composition of a tumor. Both standardized expression values and SNVs appear to be equally capable of reconstructing a similar pattern of phylogenetic relationship. This pattern is stable even when phylogenetic uncertainty is taken in account. Our results open up a new direction of somatic phylogenetics based on scRNA-seq data. Further research is required to refine and improve these approaches to capture the full picture of somatic evolutionary dynamics in cancer.

## Introduction

Phylogenetic analysis is an approach that relies on reconstructing evolutionary relationships between organisms to determine population genetics parameters such as population growth (Kingman 1982; Heled et al. 2008), structure (Müller et al. 2017a) or geographical distribution (Lemey et al. 2009; Lemey et al. 2010). Typically, the reconstructed phylogeny is not the end-goal. Using previously estimated trees, various evolutionary hypotheses can be explored, such as the evolutionary relationship of traits carried by individual taxa (Grafen et al. 1989; Pagel et al. 2004; Freckleton 2012).

Within-organism cancer evolution is increasingly being studied using population genetics approaches, including phylogenetics (Navin et al. 2011; Yuan et al. 2015; Alves et al. 2017; Schwartz et al. 2017; Caravagna et al. 2018; Singer et al. 2018; Alves et al. 2019; Caravagna et al. 2019; Detering et al. 2019; Malikic et al. 2019; Werner et al. 2019; Kuipers et al. 2020), to understand evolutionary dynamics of cancer cell populations. These approaches have shown promise to be developed into therapeutic applications in the personalized medicine framework (Gerlinger et al. 2012; Abbosh et al. 2017; Rao et al. 2020b). Specifically, the clonal composition of tumors, metastasis initiation, development, and timing can be reconstructed using phylogenetic methods (Yuan et al. 2015; Angelova et al. 2018; El-Kebir et al. 2018; Alves et al. 2019). Unlike other evolutionary processes prone to events such as hybridization or horizontal gene transfer, population dynamics of somatic cells is underpinned by a strictly bifurcating clonal process driven by cell division. This is in perfect agreement with theoretical assumptions routinely applied in stochastic phylogenetic models such as coalescent (Kingman 1982; Hudson et al. 1990; Posada 2020) or birth-death processes (Aldous 1996; Aldous 2001; Komarova 2006).

From the methodological perspective, however, cancer is an evolutionary process with unique characteristics which are not modeled in conventional phylogenetic approaches. These include a high level of genomic instability with structural changes (gene losses and duplications) which accumulate along with point mutations during the course of growth and evolution (Beerenwinkel et al. 2015; Posada 2015).

Traditional Whole Genome Sequencing (WGS) methods have been instrumental in understanding cancer mutational profiles and oncogene detection (Mardis et al. 2009; Nakagawa et al. 2018). DNA from a tissue sample is isolated and sequenced in bulk. This increases the total amount of DNA which improves coverage and reduces amplification errors. To establish the presence or absence of mutations, a variant allele frequency (VAF) is calculated and compared to a threshold, typically 10 – 20% (Strom 2016). This filters out rare mutations present only in a few reads that are likely to be false positives or sequencing errors (Petrackova et al. 2019). More recently, bulk sequencing is used to study cancer evolution using phylogenetic methods, either by comparing VAF (Zhai et al. 2017; Zhao et al. 2016; Ling et al. 2015) or estimating copy number variants (CNV) (Desper et al. 1999; Demeulemeester et al. 2016; Tarabichi et al. 2021). However, the usage of bulk sequencing in this context is problematic. Bulk samples contain cells from multiple cell lineages including non-tumor cells, such as immune or blood vessel cells (Racle et al. 2017), and there is strong evidence for a constant migration of metastatic cells between tumors (Aguirre-Ghiso 2010; Cheung et al. 2016; Reiter et al. 2017; Casasent et al. 2018). High VAF thresholds ignore tumor heterogeneity, but by lowering the threshold, mutations in non-tumor cells or clonal lineages are retained instead. Sequences or mutational profiles derived from bulk samples thus have a chimeric origin (Alves et al. 2017).

A typical assumption in classical phylogenetics is that the sequences or mutational profiles represent individual taxonomic units, either individuals or populations of closely related individuals. If these methods are used on the data from bulk samples, the reconstructed trees are not phylogenies describing an evolutionary history, but evolutionarily meaningless sample similarity trees (Alves et al. 2017). To address this issue, phylogenetic trees are reconstructed by estimating the sequential order of somatic mutations using VAF from one or multiple tumor samples (Deshwar et al. 2015; El-Kebir et al. 2018; Miura et al. 2018). Given the tumor heterogeneity and insufficient read depth to reliably estimate VAF, this is not a simple problem and the performance of current methods is limited (Miura et al. 2020).

Single-cell DNA sequencing (scDNA-seq) does not suffer from the chimeric DNA origin of bulk-sequencing as each DNA segment is barcoded to guarantee its known cell of origin. Recent progress in WGS technology made sequencing individual cells cost-efficient (Gawad et al. 2016) and this approach is now regularly used for the phylogenetic reconstruction of metastatic cancer or the subclonal structure of a single tumor (Potter et al. 2013; Roth et al. 2016; Leung et al. 2017; Myers et al. 2019). However, this increased resolution comes with additional complications. Current methods are not sensitive enough to sequence DNA from a single cell and DNA amplification is required (Gawad et al. 2016). This process suffers from a random bias with different parts of the genome amplified in different quantities or not at all (Satas et al. 2018). In addition, polymerase does not replicate DNA without error, this can have a significant impact if the replication errors occur early in the amplification process (Gawad et al. 2016). This does not only increase the error rate for identified SNVs, but a large proportion of SNVs might be simply missing (Hicks et al. 2018). The advantages associated with scDNA-seq led to the development of novel approaches that tackle these challenges using an error model to correct for amplification errors and false-positive SNV calls (Zafar et al. 2016; Zafar et al. 2018; Luquette et al. 2019; Kozlov et al. 2020).

Similar technological development led to proliferation of single-cell RNA sequencing (scRNA-seq) which, compared to traditional bulk RNA sequencing, enabled detection of gene expression profiles for individual cells in the tissue sample (Müller et al. 2017b; Olsen et al. 2018; Jerby-Arnon et al. 2018; González-Silva et al. 2020). This allows understanding tumor heterogeneity by identifying different cell populations (Andrews et al. 2018), estimating immune cell content within a tumor (Yu et al. 2019), or even identifying individual clones and subclones, as they can differ in their behavior (Fan et al. 2020). However, as the levels of RNA expression vary between genes and cells, the amplification problems of scDNA-seq that cause unequal expression and drop-out effects are more pronounced in scRNA-seq. There is an increased interest for SNV calling on scRNA-seq data using bulk-SNV callers (Chen et al. 2016; Poirion et al. 2018; Liu et al. 2019; Schnepp et al. 2019) and specialized CNV callers (Kuipers et al. 2020; Harmanci et al. 2020b; Harmanci et al. 2020a; Gao et al. 2021) as this allows for identification of mutations in actively expressed genes.

In this work, we test if expression values and SNVs inferred from scRNA-seq contain phylogenetic information to reconstruct a population history of cancer. We reconstruct phylogenetic trees using expression values and SNVs from three different datasets. We then compare and test these phylogenies against expected evolutionary history to determine whether scRNA-seq data contains phylogenetic signal.

## Methods

### Experimental design

To test if scRNA-seq data contain phylogenetic signal, we select datasets with multiple regional samples and test if cells from a regional sample are phylogenetically closer to each other than to cells from other samples.

Methods that reconstruct phylogenies from sequence data assume that the data were generated using an evolutionary process. Under this assumption, an effect of a low phylogenetic signal would produce a high mutation rate and/or star tree-like structure. However, both high mutation rate and star trees are biologically plausible in case of rapid population expansion, such as during a rapid metastatic spread (Schwarz et al. 2015). Classical methods for estimation of phylogenetic signal, such as Pagel’s *λ* (Pagel 1999) or Blomberg’s *κ* (Blomberg et al. 2003) (see Münkemüller et al. (2012) for review) are not applicable, as they test the presence of a phylogenetic signal of an observed trait based on known phylogeny. An alternative method to test whether the data does contain a phylogenetic signal that we employ in this paper is to compare the resulting phylogeny to an expected relationship, such as the expected monophyly of groups of taxa obtained from multiple regional samples, e.g. phylogenetic clusters of cells from healthy tissues or individual metastases. To further account for migration and uncertainty, we do not require cells from a single regional sample to be monophyletic but test whether cells from a regional sample are phylogenetically closer to each other than to cells from other samples using phylogenetic clustering tests.

### Dataset selection

We perform our phylogenetic analyses on three different datasets, a new UMI based dataset of breast cancer-derived xenografts (BCX), and two previously published datasets, a UMI based dataset of small intestinal neuroendocrine cancer (INC) (Rao et al. 2020a) and non-UMI based dataset of gastric cancer (GC) (Wang et al. 2021). These datasets contain primary and metastatic cells from multiple regional samples, allowing us to assess the performance of the phylogenetic analysis using the phylogenetic clustering tests. We expect that primary, metastatic, and cells from regional samples will each cluster together, forming a cell-type and region-specific clades.

The BCX dataset consisted of three individual-specific tumor samples T1, T2, and T3, seeded from the same population of cells, and two matched circulating tumor cells (CTC) samples CTC1 and CTC2. Each tumor was derived from the same population and share a common ancestor, each tumor thus represents a distinct regional sample. We expected that cells isolated from each individual and cells for each sample form clusters. As the cell lineage MDA-MB-231-LM2 is highly metastatic (Minn et al. 2005), we do not expect CTC cells to form a single monophyletic clade, but a larger number of smaller clades embedded inside the tumor cells of a sampled individual.

The INC dataset from Rao et al. (2020a) consisted of a primary tumor and a paired liver metastatic sample. Both samples contained a mixture of cancerous and non-cancerous cells (Fibroblasts, Endothelial cells, and Immune cells). We expect cancerous cells to form a cluster, with metastatic cells forming distinct clusters among the cancer cells.

The GC dataset from Wang et al. (2021) consisted of 94 cells from a primary tumor and a lymph node of three patients (GC1, GC2, and GC3). We would expect that for each patient, the lymph node cells would form a monophyletic lineage derived from the primary tumor cell, but due to the small number of cells, clustering of the primary tumor cells is also interpreted as a success.

### Preparation of the BCX dataset

MDA-MB-231-LM2 (GFP+) (Minn et al. 2005) cells were injected into the R4 mammary fat pad of Nu/J mice (250, 000 cells per mouse, 3 mice), and tumor growth was monitored for 8 weeks. Mice were euthanized when tumor size approached the endpoint (2 cm). Tumors were resected and dissociated into single cells. To extract CTC, up to 1 ml of blood was drawn immediately post euthanasia using cardiac puncture. Red blood cells were removed using RBC lysis buffer. All cells (tumor derived and circulating tumor cells) were stained with DAPI and sorted for DAPI and GFP using a BD FACSAria cell sorter. Libraries were generated using the 10x Chromium single cell gene expression system immediately after cell sorting, and sequenced on an Illumina NextSeq platform together to eliminate batch effect.

### Mapping and demultiplexing

The BCX and INC reads were mapped with the Cellranger v5.0 software to the GRCh38 v15 from the Genome Reference Consortium using the analysis-ready assembly without alternative locus scaffolds (no_alt_analysis_set) and associated GTF annotation file. The Cellranger software performs mapping, demultiplexing, cell detection, and gene quantification for the 10x Genomics scRNA-seq data.

Reads for the non-UMI GC dataset were mapped with the STAR v2.7.9a (Dobin et al. 2013), using the same reference and GTF annotation file.

### Postprocessing expression data

For the INC and BC datasets, previously published expression datasets from Rao et al. (2020a) (GSE140312) and Wang et al. (2021) (GSE158631) were used. For the BCX dataset, we have used the filtered expression values produced by the Cellranger.

#### Standardizing expression values

The filtered feature-barcode expression values from Cellranger were processed using the R Seurat v4.0.4 package (Stuart et al. 2019). Expression values from different regional samples (or individuals in case of BCX) were merged and analyzed together. The expression values for each gene were centered (*μ* = 0) and rescaled (*σ*^2^ = 1). No normalization or filtering was done at this step.

#### Discretizing expression values

To reconstruct phylogenies, expression values need to be discretized as phylogenetic software does not support tree reconstruction from continuous data. The rescaled expression values were categorized into a 5 level ordinal scale ranging from 1 (low level of expression) to 5 (high level of expression). The five-level scale was chosen to capture the data distribution of the rescaled expression values and represent a compromise between introducing data noise with too many levels or artificial similarity with only a few categories.

Interval ranges, according to which the values were categorized, were chosen according to the 60% and 90% Highest Density Intervals (HDI), the shortest intervals containing 60% or 90% of values respectively. The values inside the 60% HDI were categorized as normal, values inside the Interval ranges, according to which the values were categorized, were chosen according to the 90% HDI, but outside the 60% as increased/decreased expression and values outside the 90% HDI as a extremely increased/decreased expression.

Genes that contain only a single categorized value for each cell were removed as phylogenetically irrelevant and the discretized values were then transformed into fasta format.

#### Recording unexpressed genes as unknown data

The amount of coverage in a standard bulk RNA-seq expression analysis is usually sufficient to conclude that genes for which no molecule was detected are not expressed (Lähnemann et al. 2020). In scRNA-seq however, the sequencing coverage is very small, drop out effect is likely, and thus this assumption does not hold. This is especially a problem for non-UMI based technologies (Cao et al. 2021), but not entirely absent from the UMI-based technologies as well due to biological and technological processes (Townes et al. 2020; Hsiao et al. 2020).

According to the standard expression pipeline, these values are commonly treated as biological zeros, i.e., no detected expression of a particular gene, and have a significant impact on the data distribution during the normalization and rescaling steps (Hicks et al. 2018; Townes et al. 2020). Without an explicit model of drop out effect to account for technical or biological variation, these values might be more accurately represented as unknown values rather than true biological zeros (Van den Berge et al. 2018). We have modified the Seurat code to treat these values as unknown values (`NA` in R) and included modified functions in the `phyloRNA` package.

We will further use *data density* to describe the number of unknown values in both expression and SNV datasets, with 100% representing data set without unknown values, while 0% would represent a dataset formed entirely of unknown values.

### SNV

#### Pre-processing reads for SNV detection

The BAM files from Cellranger were processed using the Broad Institute’s Genome Analysis ToolKit (GATK) v4.2.3.0 (Poplin et al. 2018) according to GATK best practices of somatic short variant discovery. Reads were sorted, processed and recalibrated using GATK’s `SortSam`, `SplitNCigarReads` and `Recalibrate`.

#### SNV detection and filtering

To obtain SNVs for individual cells of the scRNA-seq data, first a list of SNVs were obtained by running Mutect2 (Benjamin et al. 2019), treating the data set as a pseudo-bulk sample, and retaining only the SNVs that passed all filters.

For the BCX dataset, Mutect2 was run in the tumor with matched normal sample using the parental cell linage MDA-MB-231 from Kidwell et al. (2021) and Panel of Normals derived from the same source, see supplementary materials for details.

For the IND and GC datasets, we have used Mutect2 in a tumor-only mode using the Panel of Normals and the GNOMAD germline data from the GATK best practices resource bundle.

SNVs for individual cells were then obtained by individually summarizing reads belonging to each single cell at the positions of the SNVs obtained beforehand using the pysam library, which is built on htslib (Li et al. 2009). The most common base for every cell and every position was retained, base heterogeneity and CNVs were ignored. This SNV table was then transformed into fasta format.

### Finding a well-represented subset of data

When treating the potentially unexpressed genes as unknown values, only a small proportion of the expression count values was known, with the data set derived from SNV suffering from the same problem due to the low number of reads for each cell.

While model-based phylogenetic methods can process missing data by treating the missing data as phylogenetically neutral, this significantly flattens the likelihood space which can cause artifacts, convergence problems or increase computational time (Wiens 2006; Jiang et al. 2014; Xi et al. 2016).

Published phylogenetic tools designed for single-cell DNA data sets ranged from 47 cells and 40 SNVs (Jahn et al. 2016) to 370 cells and 50 SNVs (Singer et al. 2018) or in an extreme case 18 cells and 50, 000 SNVs (Singer et al. 2018) with at most 58% of missing data across these data sets. In comparison, scRNA-seq technology can produce up to tens of thousand of cells with tens of thousand detected genes (Chen et al. 2019) and data reduction is often required.

To alleviate these issues, we employ two different filtering strategies to reduce the size of the datasets, while preserving as much information as possible, a selection strategy, where a set of high-quality cells is selected, and a stepwise filtration algorithm, where a subset of data with the highest data density is selected. Under the selection filtering strategy, a set of cells is selected, either cells of interest from the expression analysis, or a fixed subset of cells with the highest data density. This allows for a construction of datasets of specific size.

The stepwise filtering algorithm aims to find a well-represented subset of the data. By iteratively cutting out cells and genes/SNVs with the smallest number of known values, we increase the data density until a local maximum or desired data density is reached. This is equivalent to the gene/cell quality filtering during the scRNA-seq post-processing pipeline, such as in the Seurat’s standard preprocessing workflow described above, where low-quality cells and genes are removed. The advantage of this method is that a desired density can be reached with the least amount of data removed.

### Phylogenetic analysis

To reconstruct phylogenetic trees from the categorized expression values and identified SNVs, we used IQ-TREE v2.1.4 (Minh et al. 2020) and BEAST2 v2.6.3 (Bouckaert et al. 2019).

The IQ-TREE analysis was performed with an ordinal model and an ascertainment bias correction ( `-m ORDINAL+ASC` ) for the expression data, and a standard model selection was performed for the SNV data ( `-m TEST` ). Where the size of the dataset allowed, tree support was evaluated using the standard non-parametric bootstrap (Felsenstein 1985) with 100 replicates ( `-b 100` ).

The BEAST2 analysis was performed with a birth-death tree prior (Kingman 1982) with an exponential population growth (Kuhner et al. 1998), as these models most closely mimic the biological conditions of tumor growth. For the expression data, the BEAST2 was run using ordinal model available in the Morph-Models package, while the SNV values were analyzed using the Generalized Time-Reversible model (Tavaré 1986). For both the expression and the SNV data sets, BEAST2 was set to not ignore ambiguous states.

### Phylogenetic clustering tests

To test if the phylogenetic methods were able to recover expected population history, we employ Mean Pairwise Distance (MPD) (Webb 2000) and Mean Nearest Taxon Distance (MNTD) (Webb 2000). MPD is calculated as a mean distance between each pair of taxa from the same group, while MNTD is calculated as a mean distance to the nearest taxon from the same group. For each sample and samples isolated from a single individual, MPD and MNTD are calculated and compared to a null distribution obtained by permuting sample labels on a tree and calculating MPD and MNTD for these permutations. The p-value is then calculated as a rank of the observed MPD/MNTD in the null distribution normalized by the number of permutations. The MPD and MNTD are calculated using the `ses.mpd` and `ses.mntd` functions implemented in the package `picante` (Kembel et al. 2010) For the Bayesian phylogenies, MPD and MNTD were mean and 95% confidence interval. For Maximum Likelihood phylogenies, MPD and MNTD were calculated for a sample of 1000 trees from the posterior distribution and then summarized with calculated from 100 trees from the non-parametric bootstrap and summarized in the same manner as Bayesian trees.

### Code and data availability

Code required to replicate the data processing steps is available at https://github.com/bioDS/phyloRNAanalysis.

To aid in creating pipelines for phylogenetic analysis of scRNA-seq data, we have integrated a number of common tools in the R `phyloRNA` package, which is available at https://github.com/bioDS/phyloRNA.

Raw reads and expression matrices produced by Cellranger for the BCX dataset are in the NCBI GEO under the accession number GSE163210.

Alignments in the fasta format, reconstructed trees and phylogenetic clustering tests results are available at https://github.com/bioDS/phyloRNAanalysis/tree/processed_files.

## Results

### Breast cancer-derived xenografts (BCX)

#### Sample overview

In total, five samples were used in this analysis, three tumor samples (T1, T2, T3) and two CTC samples (CTC1, CTC2). The number of cells isolated from the CTC3 sample was too small for scRNA sequencing and the sample was removed from the study. The number of detected cells in the tumor samples was generally smaller than in the CTC samples, but the reverse was true for the total number of detected unique molecular identifiers (UMIs) – the number of unique mRNA transcripts (see Table 1). In the T2 sample, a large number of cells but a small number of UMIs were detected in a similar pattern to the CTC samples.

**Table 1.** An overview of the BCX dataset used in this work. In total, five samples were isolated from three individuals (Table 1a): 3 tumor samples (T1, T2, T3) and 2 circulating tumor cell samples (CTC1, CTC2). For each sample, the number of cells from fluorescent-activated cell sorting (FACS), the number of identified by Cellranger, the number of detected genes, the number of unique molecular identifiers (UMIs), UMI/Cell ratio, and the data density are reported (Table 1b).

| Sample | Cells (FACS) | Cells (Cellranger) | Genes | UMI | UMI/Cell | data density |
| --- | --- | --- | --- | --- | --- | --- |
| **T1** | 11,258 | 701 | 17k | 3,167k | 4,518 | 2.97 % |
| **T2** | 20,233 | 2,794 | 5k | 69k | 25 | 0.04 % |
| **T3** | 13,865 | 806 | 18k | 2,876k | 3,569 | 2.57 % |
| **CTC1** | 605 | 3,125 | 8k | 129k | 41 | 0.06 % |
| **CTC2** | 415 | 3,161 | 9k | 155k | 49 | 0.06 % |
| total: | 46,376 | 10,587 | 20k | 6.4M | 604 | 0.44 % |

(b) Sample overview

Compared to the fluorescent-activated cell sorting (FACS), Cellranger detected fewer cells for tumor samples, but more cells for the CTC samples. Cellranger classifies barcodes as cells based on the amount of UMI detected to distinguish real cells from a background noise (Lun et al. 2019). The large number of detected cells in the CTC samples is likely a result of lysed cells or cell-free RNA (Fleming et al. 2019). In all cases, the number of expression values across data sets was relatively low, with the best sample T3 amounting to about 3% of known expression values.

#### SNV identification

To identify SNVs in scRNA-seq data, we first identified a list of SNVs by treating the single-cell reads as a pseudo-bulk sample. The total of 21,261 SNVs that passed all quality filters were identified this way. When these SNVs were called for each individual cell, the resulting data set had data density of less than 0.13%. The expression data is expected to have higher data density than SNV because for expression quanti cation a presence or absence of a molecule is sufficient while for SNV, knowledge of each position is required. This expectation is confirmed in Table 1, where data density of the expression data is summarized. About 40% of the 10,587 cells represented in this data set did not contain any positively identified SNV after filtering out false-positives, these were relatively equally distributed among the T2 (1487), CTC1 (1379) and CTC2 (1324) samples. This represents a challenge from a data analysis perspective given the large sample size and its small data density.

#### Data reduction

With over 10,000 cells and more than 20,000 genes and SNVs, the unfiltered datasets would require substantial computational resources. An additional issue we have encountered in our data was a significant difference in the quality between individual samples, only five CTC1 and six CTC2 cells passed the quality filtering criteria of a minimum of 250 represented genes and a minimum of 500 UMI per cell, with no T2 cells passing the quality filtering. This contrasts with the T1 and T3 samples, where 701 and 806 passed the quality filtering criteria respectively. Due to this varied quality of samples, filtering data to a higher data density using the stepwise filtering algorithm leads to the removal of the low-quality samples (T1, CTC1 and CTC2), which bar us from testing the phylogenetic structure using the phylogenetic clustering tests. For this reason, we have selected a small number of cells with the least amount of missing data from each sample using the selection filtering method. The small number of cells is not sufficient to represent the full diversity of the tumor, but allow us to test the phylogenetic relationship between individual samples without introducing a bias due to an unequal size of the samples.

A total of 58 cells were retained for both the expression and SNV datasets: 20 cells for T1 and T3 samples and six cells for T2, CTC1 and CTC2 samples. In these reduced datasets, genes that were not present in any of the cells or present only in a single cell, are removed. The reduced expression data set contained 30% of known data distributed across 7,520 genes. The SNV data set contained 10% of known data distributed across 1,058 SNVs. These reduced data sets are analyzed using Maximum Likelihood and Bayesian method to further explore the topological uncertainty.

Reconstructed trees and phylogenetic tests for the data filtered to the 20%, 50% and 90% data density using the stepwise filtering algorithm are provided in the supplementary materials.

#### Phylogenetic reconstruction from expression data

The Maximum Likelihood tree reconstructed from the reduced expression data set showed significant clustering of all samples (Figure 1a). This is confirmed by the phylogenetic clustering tests where all but CTC2 cells had a significant MPD p-value (Table 2). Four out of six CTC2 cells clustered together, but on the opposite side of the tree with phylogenetic proximity to the T1 cells. This close phylogenetic relationship suggests that T1 and CTC2 were isolated from a single individual. This pattern is further reinforced as T2 cells clustered in a single compact clade with phylogenetic proximity to the CTC1 sample. When this relationship was tested with phylogenetic clustering methods, both MPD and MNTD confirmed the strong clustering signal between T2 and CTC1. The same tests were not significant for the T1-CTC2 grouping likely due to the presence of two non-clustering CTC2 cells.

**Table 2.**
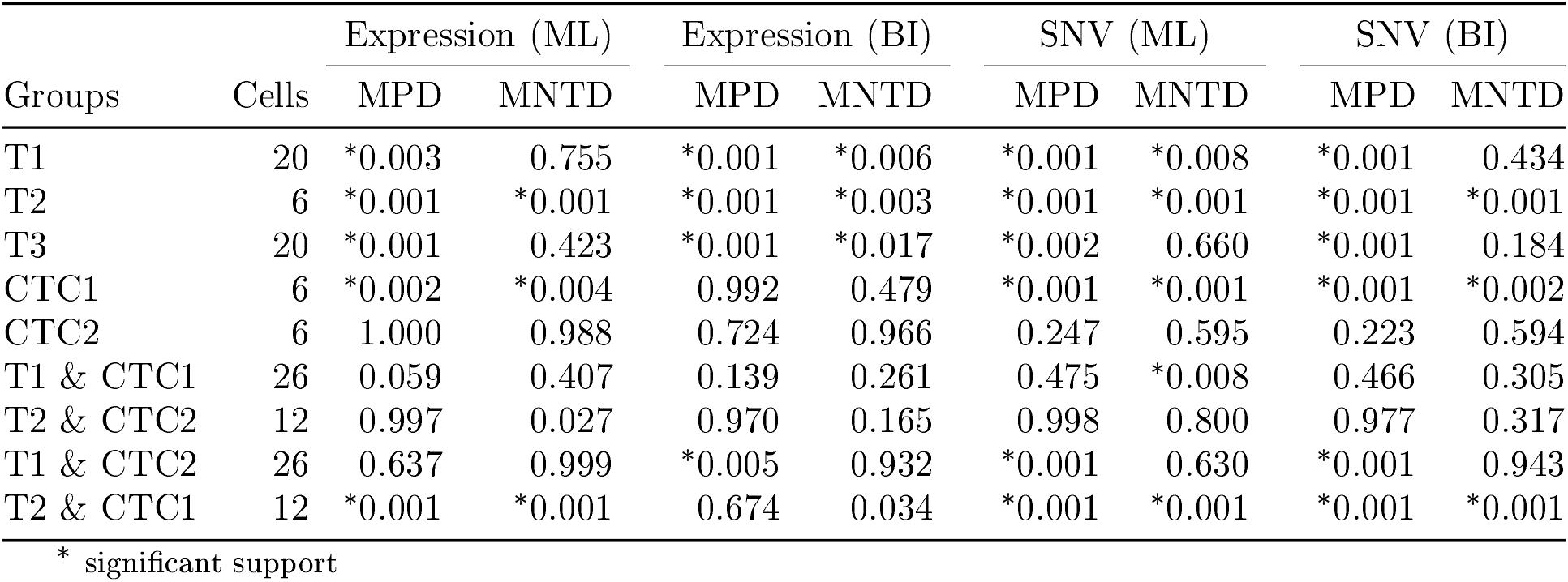
Test of phylogenetic clustering for the reduced dataset of the 58 selected cells. Mean Pairwise Distance (MPD) and Mean Nearest Taxon Distance (MNTD) calculated for the Maximum Likelihood (ML) and Bayesian (BI) trees from the expression and SNV data. P-values for MPD and MNTD were calculated for each sample (T1, T2, T3, CTC1, CTC2) and expected clustering for cells isolated from a single individual (T1 with CTC1, and T2 with CTC2) and to test a possible mislabeling between CTC1 and CTC2 samples (T1 with CTC2, and T2 with CTC1). Significant p-values at *α* = 0.05 after correcting for multiple comparisons using the False Discovery Rate method (Benjamini et al. 1995) are marked with an asterisk. Values of MNTD and MPD calculated for the Maximum Likelihood bootstrap sample and Bayesian posterior tree sample are available in the Supplementary Table 1.

| Groups | Cells | Expression (ML) |  | Expression (BI) |  | SNV (ML) |  | SNV (BI) |  |
| --- | --- | --- | --- | --- | --- | --- | --- | --- | --- |
|  |  | MPD | MNTD | MPD | MNTD | MPD | MNTD | MPD | MNTD |
| T1 | 20 | *0.003 | 0.755 | *0.001 | *0.006 | *0.001 | *0.008 | *0.001 | 0.434 |
| T2 | 6 | *0.001 | *0.001 | *0.001 | *0.003 | *0.001 | *0.001 | *0.001 | *0.001 |
| T3 | 20 | *0.001 | 0.423 | *0.001 | *0.017 | *0.002 | 0.660 | *0.001 | 0.184 |
| CTC1 | 6 | *0.002 | *0.004 | 0.992 | 0.479 | *0.001 | *0.001 | *0.001 | *0.002 |
| CTC2 | 6 | 1.000 | 0.988 | 0.724 | 0.966 | 0.247 | 0.595 | 0.223 | 0.594 |
| T1 & CTC1 | 26 | 0.059 | 0.407 | 0.139 | 0.261 | 0.475 | *0.008 | 0.466 | 0.305 |
| T2 & CTC2 | 12 | 0.997 | 0.027 | 0.970 | 0.165 | 0.998 | 0.800 | 0.977 | 0.317 |
| T1 & CTC2 | 26 | 0.637 | 0.999 | *0.005 | 0.932 | *0.001 | 0.630 | *0.001 | 0.943 |
| T2 & CTC1 | 12 | *0.001 | *0.001 | 0.674 | 0.034 | *0.001 | *0.001 | *0.001 | *0.001 |

\* significant support

**Table 3.** Test of phylogenetic clustering for the expression data when zero expression level is treated as biological zero. Mean Pairwise Distance (MPD) and Mean Nearest Taxon Distance (MNTD) calculated for the Maximum Likelihood (ML) and Bayesian (BI) trees from the expression data, with zeros treated as biological zeros. P-values for MPD and MNTD were calculated for each sample (T1, T2, T3, CTC1, CTC2) and expected clustering for cells isolated from a single individual (T1 with CTC1, and T2 with CTC2) and to test a possible mislabeling between CTC1 and CTC2 samples (T1 with CTC2, and T2 with CTC1). Significant p-values at *α* = 0.05 after correcting for multiple comparisons using the False Discovery Rate method (Benjamini et al. 1995) are marked with an asterisk.

| Groups | Cells | Expression (ML) |  | Expression (BI) |  |
| --- | --- | --- | --- | --- | --- |
|  |  | MPD | MNTD | MPD | MNTD |
| T1 | 20 | 1.000 | 1.000 | 0.998 | 0.992 |
| T2 | 6 | *0.001 | *0.001 | *0.001 | *0.001 |
| T3 | 20 | 0.543 | 0.994 | 0.804 | 0.998 |
| CTC1 | 6 | *0.001 | *0.001 | *0.001 | *0.001 |
| CTC2 | 6 | *0.004 | *0.013 | *0.001 | *0.011 |
| T1 & CTC1 | 26 | 0.992 | 0.928 | 0.968 | 0.560 |
| T2 & CTC2 | 12 | *0.001 | *0.001 | *0.001 | *0.001 |
| T1 & CTC2 | 26 | 0.997 | 0.994 | 0.991 | 0.888 |
| T2 & CTC1 | 12 | *0.001 | *0.001 | *0.001 | *0.001 |

\* significant support

**Table 4.**
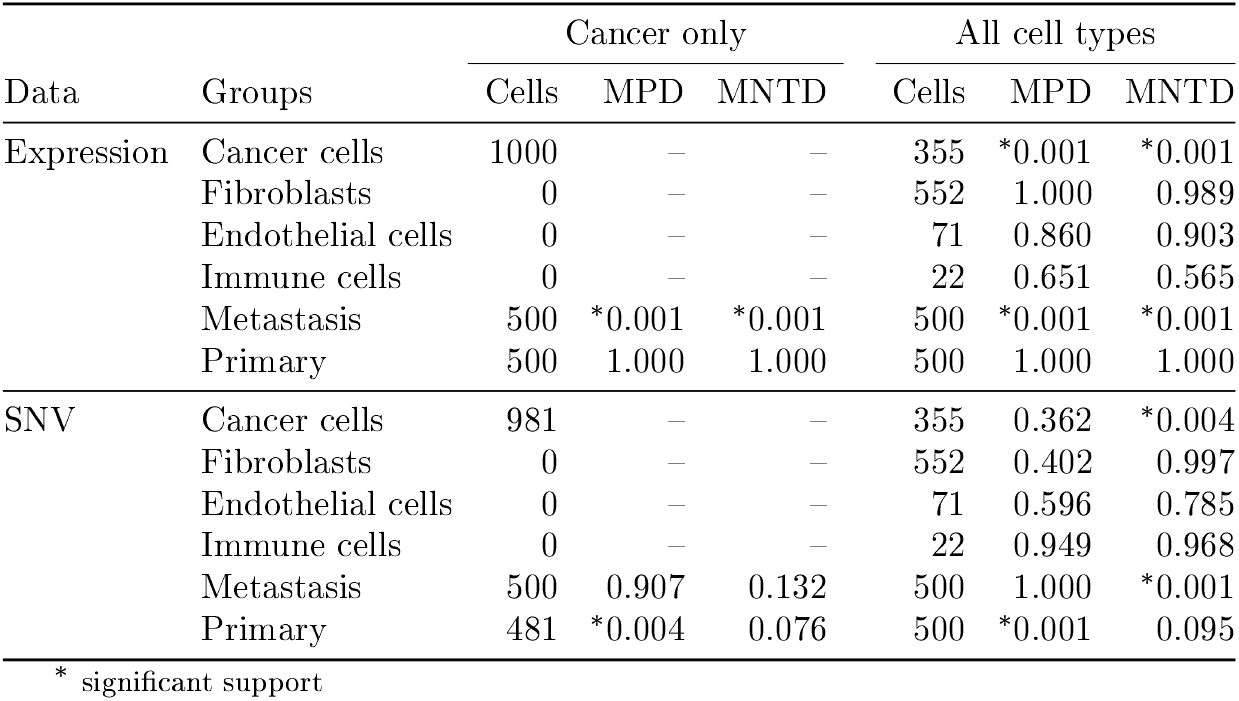
Test of phylogenetic clustering on the Maximum Likelihood trees from Rao et al. (2020a). Mean Pairwise Distance (MPD) and Mean Nearest Taxon Distance (MNTD) calculated for the phylogeny reconstructed from the dataset containing only cancer cells and from the dataset containing all cell types. P-values for MPD and MNTD were calculated for the sample of origin and cell types where applicable. Significant p-values at *α* = 0.05 after correcting for multiple comparisons using the and MNTD were calculated for the sample of origin and cell types where applicable. False Discovery Rate method (Benjamini et al. 1995) are marked with an asterisk.

**Table 5.** Test of phylogenetic clustering on the Maximum Likelihood and Bayesian trees calculated from expression and SNV data published by Wang et al. (2021). Mean Pairwise Distance (MPD) and Mean Nearest Taxon Distance (MNTD) calculated for the Maximum Likelihood and Bayesian trees reconstructed from the expression and the SNV data for patients GC1, GC2 and GC2. P-values for MPD and MNTD were calculated for the sample of origin. Significant p-values at *α* = 0.05 after correcting for multiple comparisons using the False Discovery Rate method (Benjamini et al. 1995) are marked with an asterisk.

| Data | Type | Groups | GC1 |  |  | GC2 |  |  | GC3 |  |  |
| --- | --- | --- | --- | --- | --- | --- | --- | --- | --- | --- | --- |
|  |  |  | Cells | MPD | MNTD | Cells | MPD | MNTD | Cells | MPD | MNTD |
| Expression | ML | Primary tumor | 19 | 0.171 | 0.123 | 27 | *0.001 | *0.001 | 19 | 0.829 | 0.854 |
|  |  | Lymph node | 4 | 0.830 | 0.869 | 13 | 1.000 | 1.000 | 12 | 0.138 | 0.093 |
|  | BI | Primary tumor | 19 | 0.776 | 0.995 | 27 | 0.999 | 0.953 | 19 | 0.218 | 0.232 |
|  |  | Lymph node | 4 | 0.086 | 0.035 | 13 | *0.002 | *0.001 | 12 | 0.205 | 0.333 |
| SNV | ML | Primary tumor | 19 | 0.102 | 0.111 | 27 | *0.001 | *0.001 | 19 | 0.552 | 0.231 |
|  |  | Lymph node | 4 | 0.507 | 0.092 | 13 | 1.000 | 0.945 | 12 | 0.276 | 0.208 |
|  | BI | Primary tumor | 19 | 0.117 | 0.025 | 27 | *0.001 | *0.014 | 19 | 0.531 | 0.372 |
|  |  | Lymph node | 4 | 0.955 | 0.935 | 13 | 1.000 | 0.960 | 12 | 0.487 | 0.322 |

\* significant support

**Figure 1.**
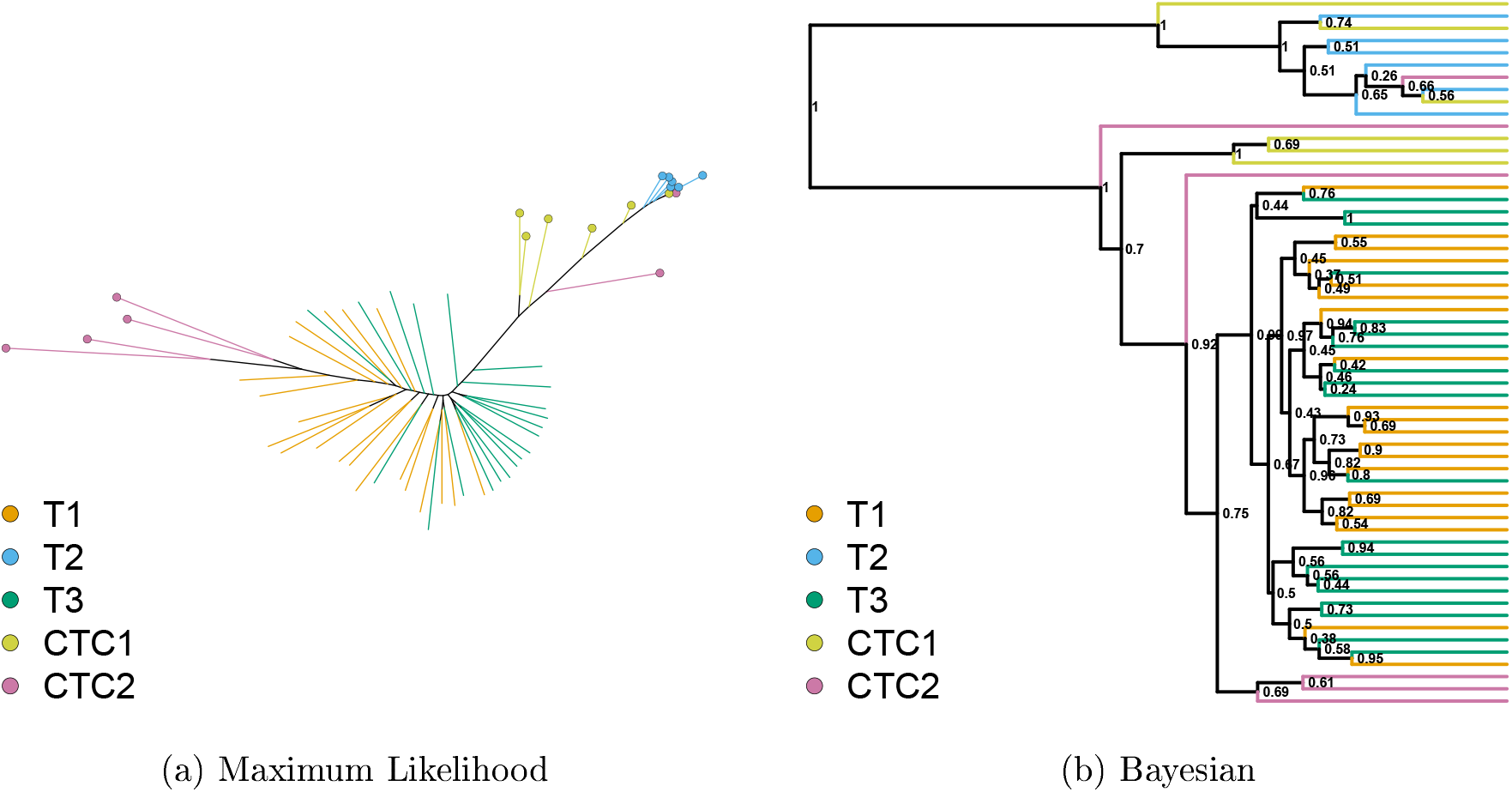
Maximum Likelihood and Bayesian trees reconstructed from the expression data for the 58 selected cells. Terminal branches are colored according to cell’s sample of origin (T1, T2, T3, CTC1, CTC2). In the Maximum Likelihood tree, the T2, CTC1 and CTC2 samples are also marked with colored circles. For the Bayesian tree, Bayesian posterior values show the topology uncertainty.

The phylogenies reconstructed from the same data using the Bayesian inference show a similar pattern of clustering (Figure 1b, Table 2), although neither CTC1 nor CTC2 formed a compact cluster. The T2 and CTC1 connection is not supported, but about half of the CTC1 cells were placed in a group with the T2 samples. Similarly to the Maximum Likelihood tree, this group was not closely related to the T1 and T3 cells, instead it formed a distantly related sister group. The relationship between T1 and CTC2 is supported by the MNTD statistics on the Bayesian phylogeny.

Neither MNTD nor MPD statistics on the Maximum Likelihood and Bayesian phylogeny supported the clustering of CTC2 cells. This might suggest that the CTC2 cells are polyphyletic, with their origin in the seeding population before the injection. This is not unlikely given that the cell lineage used (MDA-MB-231-LM2) is highly metastatic (Minn et al. 2005).

In addition to testing on the best phylogeny, we have integrated the topological uncertainty of the reconstructed phylogenies by performing the phylogenetic clustering tests on the 100 bootstrap replicates from the Maximum Likelihood analysis and a sample of 1000 trees from the Bayesian posterior tree sample. The distribution of MPD and MNTD p-values calculated on each tree were then summarized using mean and 95% confidence interval. The majority of relationships from the best tree were also supported by the tree samples (Supplementary Table 1). This suggests that while there is high uncertainty in the data and reconstructed topologies, we can reconstruct broad topological patterns with relatively high certainty.

#### Phylogenetic reconstruction from the SNV data

The Maximum Likelihood tree reconstructed from the reduced SNV dataset (Figure 2) displayed similar but weaker patterns to the one reconstructed from the expression data. The CTC2 cells no longer formed two compact clusters and were dispersed along the tree. Similarly to the expression data, the T2 and CTC1 cells were placed together on a long branch suggesting a long shared evolutionary history. However, unlike the expression data, the T1 and T3 were more interspersed with very short branches. The phylogenetic clustering tests confirm the grouping of all samples (Table 2), except for the CTC2 sample, in addition to the putative relationship between T1 and CTC2, and T2 and CTC1 samples. This reinforces the hypothesis about possible mislabeling between CTC1 and CTC2 samples.

**Figure 2.**
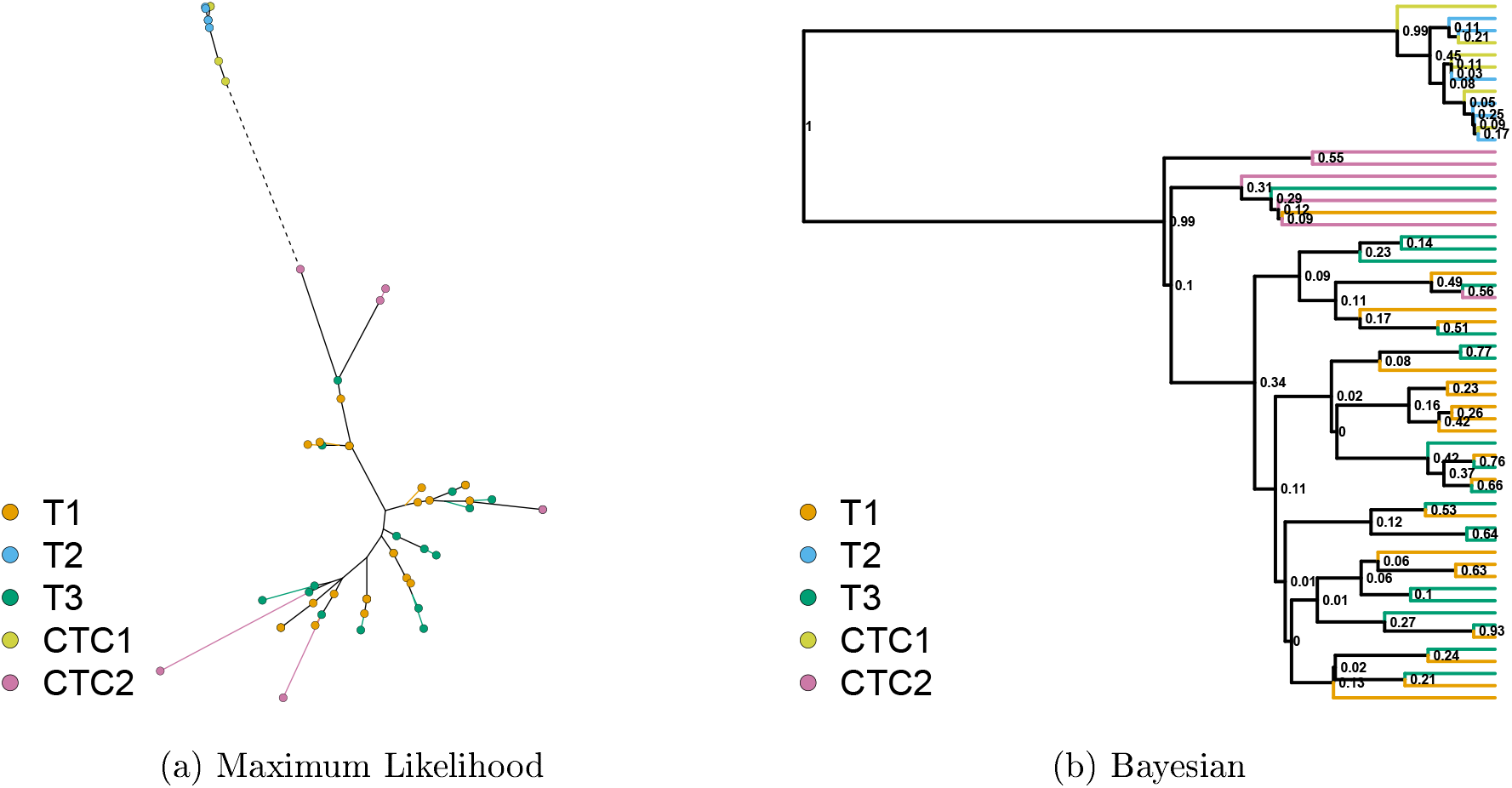
Maximum Likelihood and Bayesian trees reconstructed from the SNV data for the 58 selected cells. Terminal branches are colored according to cell’s sample of origin (T1, T2, T3, CTC1, CTC2). In the Maximum Likelihood tree, cells are also marked with colored circles and an extremely long branch leading to T2 cells (dashed line) was collapsed. For the Bayesian tree, Bayesian posterior values show the topology uncertainty.

A similar pattern of sample clustering can be observed on the Bayesian phylogeny reconstructed from the same data (Figure 2b), with T2 and CTC1 cells placed on a distantly related sister branch to all other samples. The T1 and T3 cells are still interspersed, but the CTC2 cells seem to cluster together more closely. Like with the expression analysis, when these relationships are stable when the topological uncertainty is integrated into the phylogenetic clustering tests (Supplementary Table 1).

Additional topological comparison between trees from SNV and expression data can be found in supplementary materials.

#### Biological zero or unknown value

To test the assumption if the zero expression values should be treated as unknown data rather than biological zeros, i.e., no expression of a particular gene, we have reconstructed the phylogenies from the scRNA-seq expression by treating the zeros in the dataset as biological zeros. Data were processed as per the standard methodology to get the alignments, but instead of treating the zeros as an unknown position, they were treated as a category 0 in addition to the 5 level ordinal scale. Phylogenies were then reconstructed using both Maximum Likelihood and Bayesian methods with sample clustering explored using the phylogenetic clustering tests.

In the phylogenies reconstructed from the expression data when zero is treated as a biological zero (Figure 3), the CTC2 cells did not form a cluster but clustered closely with the T1 and CTC2 cluster. This cluster was no longer placed as a sister branch to the T1 and T3 cells but was deeply nested. The T1 and T3 samples were less interspersed than when zero is treated as unknown data. This change in the phylogenetic structure is supported by the phylogenetic clustering tests, with T1 and T3 no longer being supported and instead, the clustering of CTC2 cells is being supported in both the Maximum Likelihood and Bayesian phylogenies. Likewise, the T1 and CTC2 grouping is not supported, as the CTC2 cells group together with the CTC1 and T2 samples.

**Figure 3.**
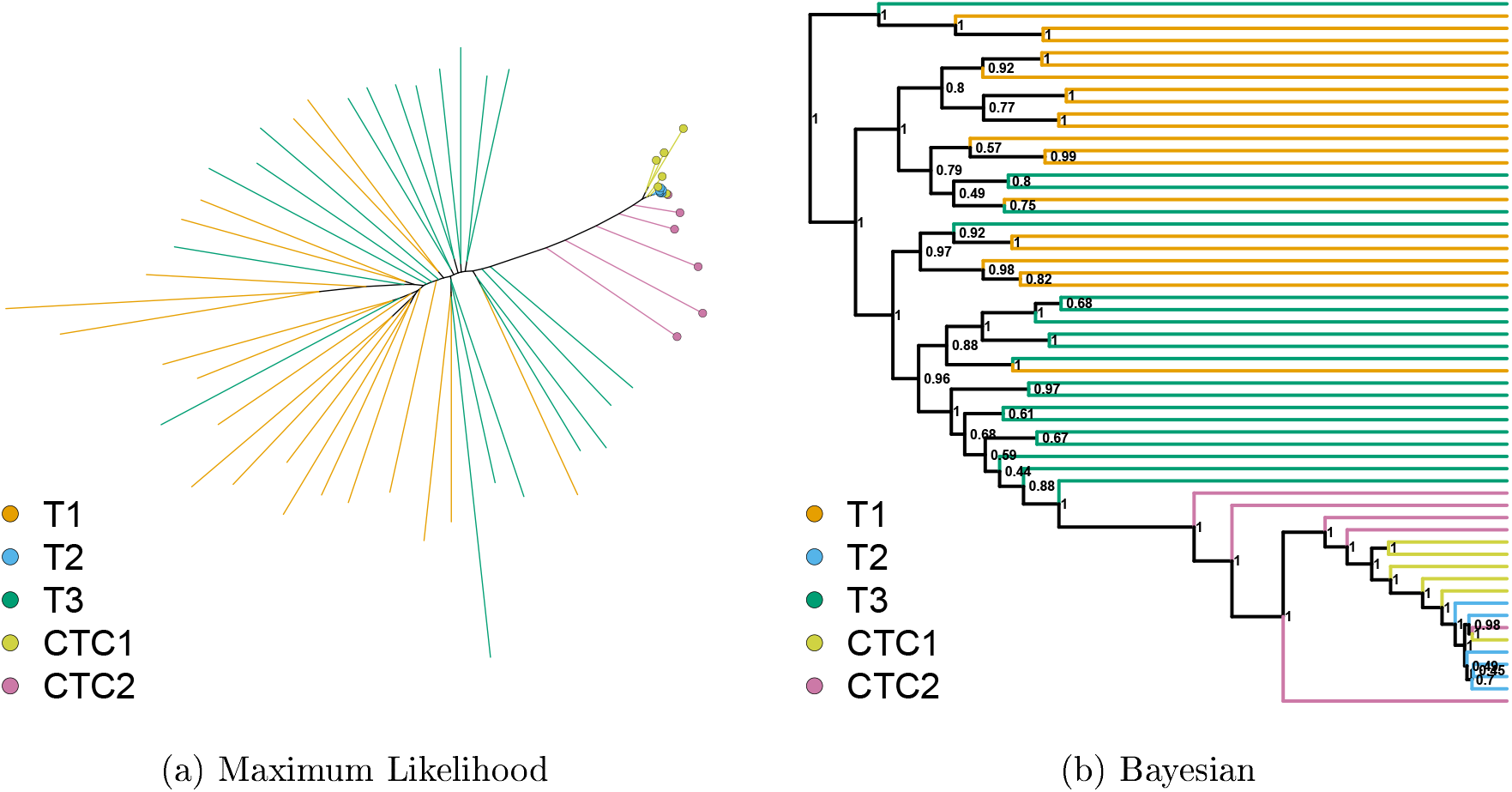
Maximum Likelihood and Bayesian trees reconstructed from the expression data for the 58 selected cells. Terminal branches are colored according to cell’s sample of origin (T1, T2, T3, CTC1, CTC2). In the Maximum Likelihood tree, the T2, CTC1 and CTC2 samples are also marked with colored circles. For the Bayesian tree, Bayesian posterior values show the topology uncertainty.

These results do not provide a conclusive answer on which assumption should be preferred. Assuming all zeros to be biological zeros will bias the model as many of those might be technical zeros instead. At the same time, the pattern of expression and non-expression seems to carry information. This information is lost when all zeros are assumed to be technical zeros and thus unknown data. For our datasets, the assumption of zeros as technical zeros and thus unknown data seems to create better agreement in the phylogenetic structure between the expression and SNVs and thus should be preferred. However, our datasets also suffered from unequal data quality issues (Table 1), and under different conditions, assuming zeros as biological zeros might be preferred.

### Intestinal neuroendocrine cancer (INC)

Cells were labeled according to their sample of origin (primary tumor and metastasis) and their cell type, which was determined by replicating the analysis from Rao et al. (2020a). We have derived two subsets from the expression and SNV data for the INC dataset from Rao et al. (2020a), a subset with all cell types and a subset with cancer cells only. To do this, cells were labeled according to their sample of origin (primary tumor and metastasis) and their cell type, which was determined by replicating the analysis from Rao et al. (2020a). For each subset, 1000 cells with the least amount of missing data were selected, 500 from the primary tumor and 500 from the metastatic sample. However, not all cells found in the expression subsets were found in the SNV data. This is likely due to a different version of the Cellranger software used in this work compared to the Rao et al. (2020a). In both derived subsets from the expression data, metastatic cells showed a strong clustering tendency (*p* = 0.001) into several large clades (Figure 4). This suggests a strong phylogenetic relationship with several well-preserved lineages. In addition, in the derived subset containing all cell types, the cancer cells showed a significant clustering (*p* = 0.001), while other cell types showed the opposite tendency (Figure 4). However, In addition, in the derived subset containing all cell types, the cancer cells showed a significant the cancer clade contained deeply nested clades of Endothelial cells and Immune cells. A similar albeit significantly weaker pattern of cancer cell clustering can be observed on the trees derived from the SNV data (Figure 4, Figure 4). In both subsets derived from the SNV data, the primary cells clustered together, but the pattern was less consistent and confirmed only by one of the two tested statistics.

**Figure 4.**
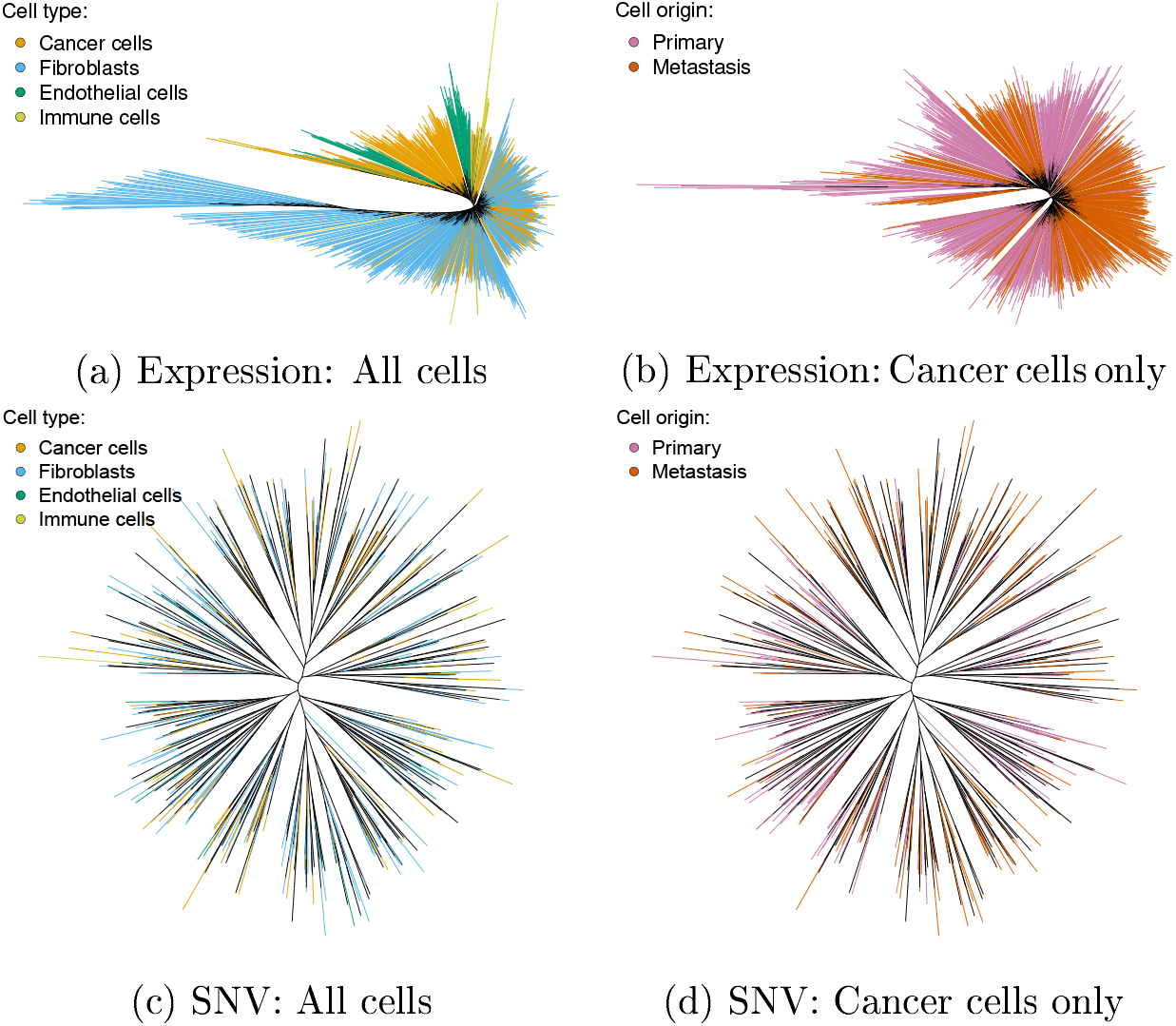
Maximum Likelihood trees constructed from the expression and SNV data published by Rao et al. (2020a). Terminal branches are colored according to cell’s type or sample of origin. In the tree reconstructed from expression data for all cells (Figure 4a), the vast majority of cancer cells cluster in a single clade. The tree reconstructed from expression data for cancer cells only (Figure 4b) shows a strong clustering of primary and metastatic cells. While the metastatic cells are not clustered in a single clade, multiple metastatic events are biologically plausible. In the trees reconstructed from the SNV data (Figure 4c, Figure 4d), primary and metastatic cells, as well as cells of different type, are relatively evenly distributed without any apparent clustering.

### Gastric cancer (GC)

For both the expression and SNV data from the gastric cancer dataset published by Wang et al. (2021), only a single patient showed significant clustering of lymph nodes (Figure 5). Poor separation of primary and lymph node cells from the expression levels was pointed out in the original study (Wang et al. 2021). Additionally, non-UMI based methods suffer from an increased error rate through zero-count in ation (Cao et al. 2021) and amplification variability (Townes et al. 2020). In the absence of a strong phylogenetic signal shared by a large percentage of genes, this additional noise is making a phylogenetic reconstruction difficult, if not impossible. At the same time, the typically higher coverage in the non-UMI based sequencing compared to the UMI should improve the identification of SNVs and decrease the misspecification error. This might suggest that different strategies for the phylogenetic reconstruction should be applied to UMI and non-UMI based sequencing.

**Figure 5.**
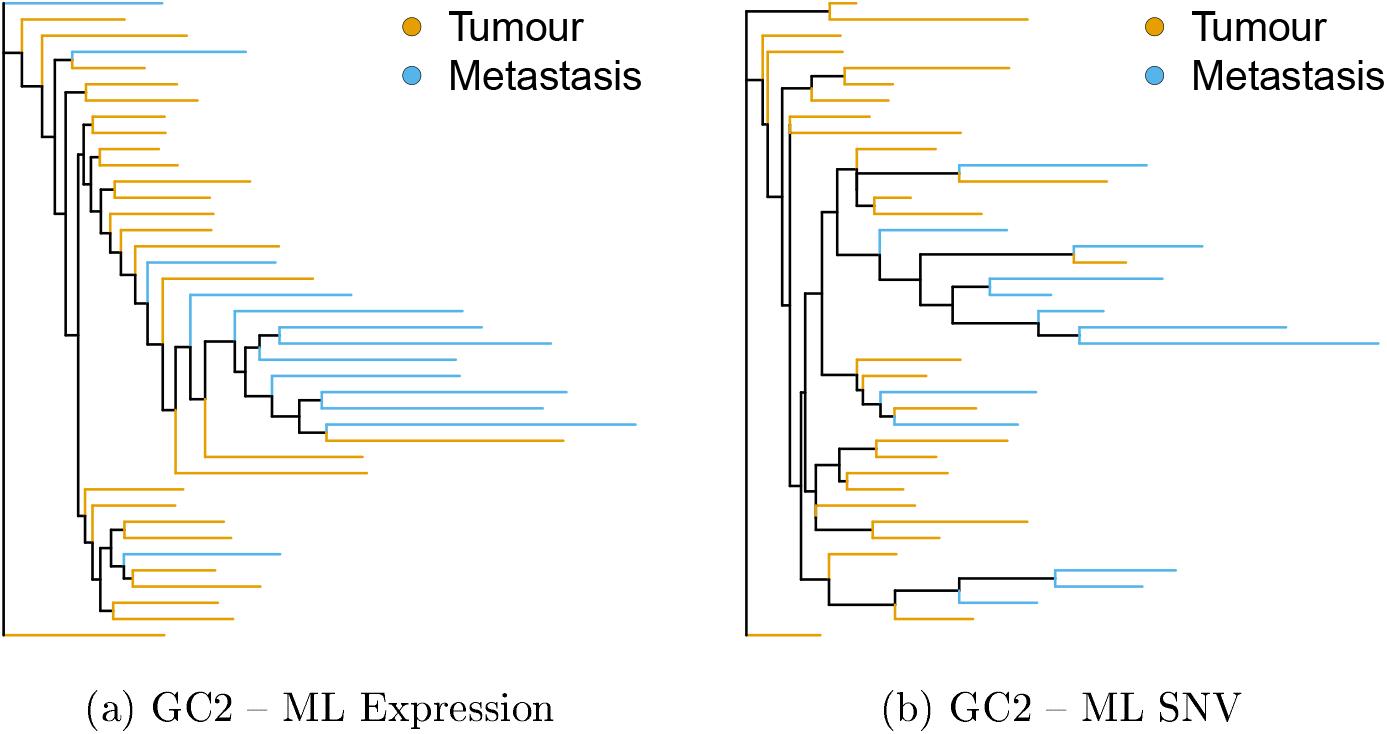
Maximum Likelihood trees for the patient G2 constructed from the expression and SNV data published by Wang et al. (2021). Terminal branches are colored according to cell’s sample of origin. Only the patient G2 shows a significant clustering signal both on the trees from Expression and SNV data. For all trees, see Supplementary Figure 3 and Supplementary Figure 4

## Discussion

Phylogenetic methods using scDNA-seq data are becoming increasingly common in tumor evolution studies. scRNA-seq is currently used for studying expression profiles of cancer cells and their behavior. However, while clustering approaches to identify cells with similar expression profiles are common and frequently used, scRNA-seq data are yet to be used in phylogenetic analyses to reconstruct the population history of somatic cells. To test if the scRNA-seq contains a phylogenetic signal to reliably reconstruct the population history of cancer, we have performed an experiment to produce a known history by infecting immunosuppressed mice with human cancer cells derived from the same population. Then using two different forms of scRNA-seq data, expression values, and SNVs, we reconstructed phylogenies using Maximum Likelihood and Bayesian phylogenetic methods. By comparing the reconstructed trees to the known population history, we confirmed that scRNA-seq contains a phylogenetic signal to reconstruct the population history of cancer, with both the expression values and SNVs producing a similar phylogenetic pattern. However, this signal is burdened by uncertainty in both the source data as well as reconstructed phylogeny. Accurate phylogenies might thus need an explicit error model to account for this increased uncertainty (Hicks et al. 2018). Still, by taking this topological uncertainty into account, we can make a conclusion about the structural relationship of individual cells. This highlights that scRNA-seq can be utilized to explore both the physiological behavior of cancer cells and their population history using a single source of data.

While the expression phylogeny can be obtained for virtually no added cost, the choice of normalization method, batch-effect correction, discretization, or the choice between biological and technical zeros, can greatly influence the phylogeny. The phylogenies reconstructed from SNV do not suffer from these decisions and SNV-calling pipelines will likely differ only in the number of false negatives and false positives.

Without any specialized phylogenetic or error models for the scRNA-seq data, conventional methods and software tools developed for systematic biology are able to reconstruct population history from this data, potentially at low computational cost. This implies that more accurate inference will be possible when and if specialized models and software are developed, and serious computational resources are employed. For example, computationally more intensive standard non-parametric bootstrap or Bayesian methods on the unfiltered data sets are certainly within the reach of modern computing clusters. This is a future direction for research.

In this work, we tested for phylogenetic signal on three data sets, a new data set consisting of 5 tumor samples seeded using a population sample, and two previously published data sets consisting of a primary tumor with a paired lymph node or a metastatic samples. Due to the nature of the experiment and the amount of uncertainty in the scRNA-seq data, this barred us from a more detailed exploration of the tree topology as only broad patterns, the phylogenetic clustering of cells according to sample and individual of origin, could be considered. Our clustering analyses show that the phylogenetic trees conform broadly to the expected shapes under different experimental conditions, and thus that expression and SNV data can both be used to infer phylogenetic trees from scRNA-seq. Nevertheless, our results also demonstrate that all such trees contain significant uncertainty, so new datasets and methods will be required to extend this work.

The degree to which low and uneven gene expression plays a role in scRNA-seq requires special attention, especially for non-UMI based data sets, as this causes not only a large proportion of missing data, but also burdens the known values with a significant error rate. Research should aim at trying to quantify this expression-specific error rate and build specialized models to include the uncertainty about the observed data in the phylogenetic reconstruction itself. This could potentially include removing a large proportion of low-coverage data in favor of robust analysis and proper uncertainty estimation of the inferred topology.

The estimation of the topological uncertainty, be it the Bootstrap branch support or the Bayesian posterior clade probabilities, is a staple for phylogenetic analyses. Currently existing methods for the phylogenetic analysis of scDNA-seq, such as SCITE (Jahn et al. 2016), SiFit (Zafar et al. 2017), or SCIΦ (Singer et al. 2018), do not provide this uncertainty estimate. This makes interpretation of the estimated topology difficult because a single topology can only be marginally more accurate than a number of alternative topologies. Of packages we are aware of, only CellPhy, through its integration in the phylogenetic software RAxML-NG (Kozlov et al. 2019), provides an estimate of topological uncertainty through the bootstrap method. Bayesian methods could be a solution as they provide an uncertainty estimate through the posterior distribution. However, they are significantly more computationally intensive than Maximum Likelihood methods. Instead, as the size of single-cell data sets will only increase, bootstrap approximations optimized for a large amount of missing data need to be developed to provide a fast and accurate estimate of topological uncertainty.

An aspect of scRNA-seq expression data that was not considered here is correlated gene expression (Wang et al. 2004; Bageritz et al. 2019). A single somatic mutation could thus induce a change of expression of multiple genes. This might be problematic given that phylogenetic methods assume that individual sites are independent and this would cause an overestimation of a mutation rate. However, phylogenetic methods are generally rather robust to a wide range of model violations (Huelsenbeck 1995a; Huelsenbeck 1995b; Song et al. 2010; Philippe et al. 2011). In addition, by randomly sampling sites, the bootstrap analysis does explore solutions that would arise from this model violation. An investigation of the effect of correlated gene expression on the estimated phylogeny provides an interesting direction for further research.

Multiomic approaches are increasingly popular as they integrate information from multiple biological layers (Bock et al. 2016; Hasin et al. 2017; Nam et al. 2020). While CNVs were ignored in this paper, it is possible to detect large-scale CNVs from scRNA-seq data (Müller et al. 2018; Kuipers et al. 2020; Harmanci et al. 2020b; Harmanci et al. 2020a; Gao et al. 2021). Combined with the SNVs and expression data as analyzed in this paper, this enables a multiomic approach using just a single scRNA-seq data source, without the additional cost of DNA sequencing.

## Acknowledgement

We thank Dr. Jon Preall and the Genomics Technology Development Core (CSHL) for scRNA-seq library preparation, and Pamela Moody and the Flow Cytometry Facility (CSHL) for support with single-cell sorting. We acknowledge Suzanne Russo for technical assistance with animal experiments.

AG and JCM acknowledge support from the Royal Society te Apārangi through a Rutherford Discovery Fellowship (RDF-UOC1702), AG, JCM, RL, and SDD acknowledge support of from an Endeavour Smart Ideas grant (UOOX1912), AG acknowledges support from a Data Science Programmes grant (UOAX1932), SDD acknowledges support from a Rutherford Discovery Fellowship (RDF-UOO1802) and the NHI/NCI grant (1K99CA215362-01), and DLS acknowledges support from the NCI grant (5P01CA013106-Project 3). We would also like to acknowledge the CSNL Next-Gen Sequencing Core (NCI-2P30CA45508).

## Supplementary materials

### Constructing the Panel of Normals and normal samples from the MDA-MB-231 cell lineage

The data from the Kidwell et al. (2021) contain a mixture of macrophages and MDA-MB-231 cells with and without mitochondrial transfer. To create the Panel of Normals, we have used all reads, but for the matched normal samples, only reads belonging to the MDA-MB-231 cells without mitochondrial transfer were used.

The fastq files were downloaded from the NCBI GEO database (ascension number GSE181410), mapped using the Cellranger and preprocessed using the GATK best practices as per the methodology section. The Panel of Normals was then constructed as per GATK instructions by first running Mutect2 in a tumor-only mode, merging the resulting variants into a database and then creating a Panel of Normals variant file from this database.

For the normal samples, we have used the preprocessed bam files from the previous steps. Only the bam files that contained MDA-MB-231 cells without mitochondrial transfer were retained (GSM5501832 and GSM5501833). To remove macrophages, the bam files were then filtered using the cell barcodes from the h5-Seurat expression analysis, that identified MDA-MB-231 cells without mitochondrial transfer (samples 2A and 2B in the Seurat object). These filtered bam files were used as matched normal samples in the main analysis.

### Integrating topological uncertainty

To investigate how the topological uncertainty influences the phylogenetic relationship between samples, we perform the phylogenetic clustering tests over 100 bootstrap samples of the Maximum Likelihood trees and over sample of 1000 posterior trees from the Bayesian inference. First we have sub-sampled the posterior tree sample to gain a sample of 1000 trees using the `logcombiner` from the BEAST2 package. Then, we calculate the MPD and MNTD p-values for each tree in the Maximum Likelihood and Bayesian tree sample. Finally, we summarize each sample using the mean and the 95% confidence interval. The calculated values for the Maximum Likelihood and Bayesian trees reconstructed from the subset of 58 cells from the expression and SNV data are presented in Supplementary Table 1. The majority of relationship that were present on the best tree is stable when the topological uncertainty is taken in account. MNTD is less stable than MPD, likely due to a higher sensitivity of MNTD to patterns closer to the tips of a tree.

### Comparison of topologies obtained from different methods

To further explore how the topologies reconstructed from SNV and expression data differ, we have calculated pairwise distances of the best trees from SNV and Expression data obtained from Maximum Likelihood and Bayesian inference. To calculate a topological distance between two trees, we chose the normalized Matching Split Information Distance (nMSID) (Smith 2020a) from the package TreeDist (Smith 2020b), which is a generalization of Robinson-Fould distance that calculates the amount of shared information between two trees and the RNNI distance (Gavryushkin et al. 2018) calculated using the FINDPATH algorithm (Collienne et al. 2021). The RNNI distance is a distance in the RNNI space (Gavryushkin et al. 2018), which is defined only for ranked topologies and thus can be calculated only for ultrametric trees produced in this paper by the Bayesian Inference. One advantage of the RNNI distance for our purposes is that the expected distance between two random trees is known and fixed at 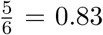 of diameter (maximum distance) (Collienne 2021), while for advantage of the RNNI distance for our purposes is that the expected distance between two random the nMSID, the expected random distance needs to be simulated. The pairwise distances between trees reconstructed from SNV, expression values, and expression values with not expressed genes treated as biological zeros (Expr0), calculated using Maximum Likelihood and Bayesian inference are summarized in Supplementary Table 2, together with distances to their tree samples. Trees reconstructed from SNV and expression values are relatively distant (nMSID= 40 – 44), but so are trees from expression data reconstructed by different methods (nMSID= 0.37). Curiously, when not expressed genes are treated as biological instead of technical zeros, the reconstructed trees are closer under the nMSID to both Expression and SNVs phylogenies from either method. An interesting pattern also emerges in the RNNI space. While the SNV tree seems to be relatively equidistant from both expression trees (RNNI= 0.59 and 0.56 for Expr and Expr0 respectively), the expression trees are significantly further away from each other (RNNI= 0.71). In the case of SNVs, the best tree was also quite distant from its bootstrap or posterior sample of trees (mean nMSID=0.33 and 0.22 for ML and BI respectively), this is in contrast with the expression trees, where their sample similar than expected by random chance, with an expected random distance of 0.54 for nMSID and was densely spaced around the best tree. For both nMSID and RNNI, all examined trees were more 0.83 for RNNI. This again highlights the amount of uncertainty that is in the estimate and the importance of phylogenetic clustering tests to recover the required pattern, which was relatively consistent for all trees.

**Supplementary Table 1.** Test of phylogenetic clustering for the reduced dataset of the 58 selected cells. Mean Pairwise Distance (MPD) and Mean Nearest Taxon Distance (MNTD) calculated for the Maximum Likelihood (ML) and Bayesian (BI) trees from the expression and SNV data. P-values for MPD and MNTD were calculated for each sample (T1, T2, T3, CTC1, CTC2) and expected clustering for cells isolated from a single individual (T1 with CTC1, and T2 with CTC2) and to test a possible mislabeling between CTC1 and CTC2 samples (T1 with CTC2, and T2 with CTC1). P-values were calculated for the sample of 100 bootstrap trees and 1000 posterior trees from the Maximum Likelihood and Bayesian analyses respectively, and this distribution of p-values is summarized with mean and 95% confidence interval. Significant p-values at *α* = 0.05 after correcting for multiple comparisons using the False Discovery Rate method (Benjamini et al. 1995) are marked with an asterisk.

| Groups | Cells | Expression (ML) |  | Expression (BI) |  | SNV (ML) |  | SNV (BI) |  |
| --- | --- | --- | --- | --- | --- | --- | --- | --- | --- |
|  |  | MPD | MNTD | MPD | MNTD | MPD | MNTD | MPD | MNTD |
| T1 | 20 | *0.001 (0.001–0.003) | 0.752 (0.637–0.877) | *0.001 (0.001–0.001) | *0.007 (0.001–0.019) | *0.001 (0.001–0.002) | 0.059 (0.001–0.192) | *0.001 (0.001–0.001) | 0.167 (0.001–0.563) |
| T2 | 6 | *0.001 (0.001–0.001) | *0.001 (0.001–0.001) | *0.002 (0.001–0.006) | *0.004 (0.001–0.017) | 0.017 (0.001–0.001) | 0.032 (0.001–0.005) | *0.003 (0.001–0.001) | *0.005 (0.001–0.007) |
| T3 | 20 | *0.001 (0.001–0.001) | 0.392 (0.280–0.526) | *0.001 (0.001–0.002) | 0.040 (0.002–0.095) | *0.002 (0.001–0.004) | 0.386 (0.021–0.780) | *0.001 (0.001–0.002) | 0.268 (0.009–0.703) |
| CTC1 | 6 | *0.004 (0.001–0.012) | *0.005 (0.001–0.013) | 0.988 (0.971–0.999) | 0.470 (0.280–0.621) | *0.011 (0.001–0.002) | 0.024 (0.001–0.015) | *0.002 (0.001–0.002) | *0.007 (0.001–0.023) |
| CTC2 | 6 | 1.000 (1.000–1.000) | 0.987 (0.937–1.000) | 0.701 (0.648–0.747) | 0.946 (0.845–1.000) | 0.237 (0.161–0.347) | 0.491 (0.230–0.727) | 0.175 (0.083–0.247) | 0.433 (0.183–0.628) |
| T1 & CTC1 | 26 | 0.051 (0.022–0.082) | 0.427 (0.273–0.641) | 0.146 (0.110–0.180) | 0.281 (0.049–0.499) | 0.426 (0.245–0.557) | 0.084 (0.001–0.257) | 0.393 (0.262–0.521) | 0.225 (0.001–0.748) |
| T2 & CTC2 | 12 | 0.995 (0.986–1.000) | 0.092 (0.002–0.321) | 0.949 (0.912–0.979) | 0.127 (0.004–0.304) | 0.950 (0.663–1.000) | 0.539 (0.119–0.854) | 0.938 (0.859–0.996) | 0.379 (0.077–0.680) |
| T1 & CTC2 | 26 | 0.574 (0.360–0.716) | 1.000 (0.998–1.000) | *0.005 (0.002–0.009) | 0.889 (0.741–0.988) | *0.001 (0.001–0.002) | 0.450 (0.062–0.826) | *0.001 (0.001–0.003) | 0.465 (0.059–0.896) |
| T2 & CTC1 | 12 | *0.001 (0.001–0.001) | *0.001 (0.001–0.001) | 0.654 (0.556–0.758) | 0.053 (0.001–0.173) | *0.004 (0.001–0.027) | 0.060 (0.001–0.723) | *0.001 (0.001–0.001) | *0.005 (0.001–0.003) |

\* significant support

**Supplementary Table 2.**
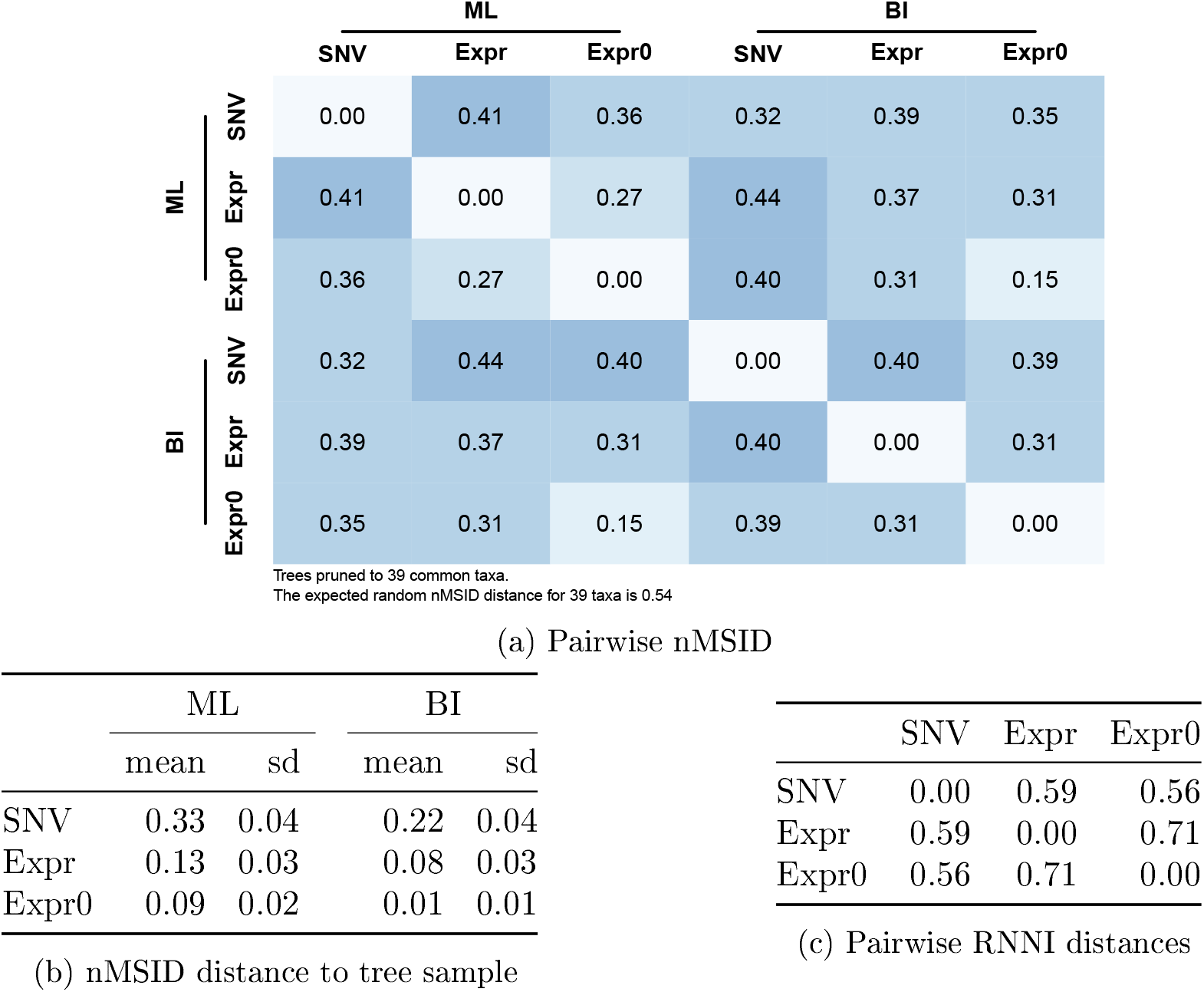
Comparison of distances between trees made from SNV and expression data from the BCX dataset. Table 2a shows pairwise distances between the best trees from SNV, expression values, and expression values where not expressed genes are considered true biological zeroes (Expr0), from both Maximum Likelihood and Bayesian Inference. Table 2b summarizes the distance between these trees and their respective bootstrap or posterior tree samples. Table 2c shows pairwise RNNI distances between the best trees from SNV, expression values, and expression values where not expressed genes are considered true biological zeroes from Bayesian Inference

### Data reduction using the stepwise filtering algorithm

Using the stepwise filtering algorithm, we iteratively remove cells and genes/SNVs with the smallest number of known values, until a desired data density is reached. By using this method, the least amount of data is removed. Here we investigate the effect of unknown data on the reconstructed topology by preparing datasets with different data densities using our stepwise filtering algorithm.

The expression data set filtered to 20% density contained 1,627 cells and 6,187 genes. Cells were mainly represented by T1 and T3 samples which form over 92% of the data set. In contrast, other in an unfiltered data set. In filtering to 50% density, the numbers decreased to 1,454 cells and 1,634 genes. The sample diversity also decreased, with T2 dropping out entirely. When filtered to 90% samples (T2, CTC1, CTC2) were significantly underrepresented despite their larger amount of cells density, the data set was reduced to 593 cells and 528 genes. The data diversity further decreased to T1, T3 and CTC2, with the CTC2 sample reduced to 2 cells.

When the SNV data set was filtered to 20% density, the numbers decreased significantly − to 870 cells and 317 SNVs, with only T1, T3 and CTC2 samples present. In subsequent filtering to 50% data density, the dataset was reduced to 254 cells and only 69 SNVs, with subsequent filtering to the 90% reducing the dataset further to 60 cells and 8 SNVs.

We inferred Maximum Likelihood trees of the expression data filtered to 20%, 50%, and 90% data density (Supplementary Figure 1). In the reconstructed phylogeny at the 20% density filtering (Supplementary Figure 1a), individual tumor samples did not form three separate clades, but a large number of smaller clades. These clades are distributed along the central spine of the unrooted maximum likelihood tree and have little internal structure. The T2, CTC1, and CTC2 samples form relatively compact clades, while the more represented T1 and T3 clades are generally intermixed. The MNT and MNTD test confirm this (Supplementary Table 3), with T2, CTC1, and CTC2 showing significant clustering signal. When the data is filtered to 50% and 90% data density (Supplementary Figure 1), the previously significant relationship disappears, likely due to the reduction of T2, CTC1 and CTC2 samples into a small number of cells.

In the trees reconstructed from the SNV data, the CTC2 cells do cluster (Supplementary Figure 2), but this relationship is not significant (Supplementary Table 3), likely due to the small number of cells remaining compared to a large number of cells from the T1 and T3 datasets. Only the T1 and the putative relationship between T1 and CTC2 cells was supported, but this support disappeared when the data was further filtered. Due to the small number of SNVs for the dataset filtered to 90% data density, many cells were identical, with small or collapsed branches (Supplementary Figure 2c).

The data filtered with the stepwise filtering algorithm failed to show a convincing clustering signal for each sample. This is likely caused by the variable quality of our sample and thus the results should be interpreted in this context. On a dataset that is not burdened by similar issues, the stepwise filtering algorithm might be the preferred method. The partial clustering signal for the lowest density filtering shows that the phylogenetic methods can handle well large amount of missing data. This means that data reduction should be performed carefully to limit the required amount of computational burned and n to reduce the amount of missing data.

**Supplementary Figure 1.**
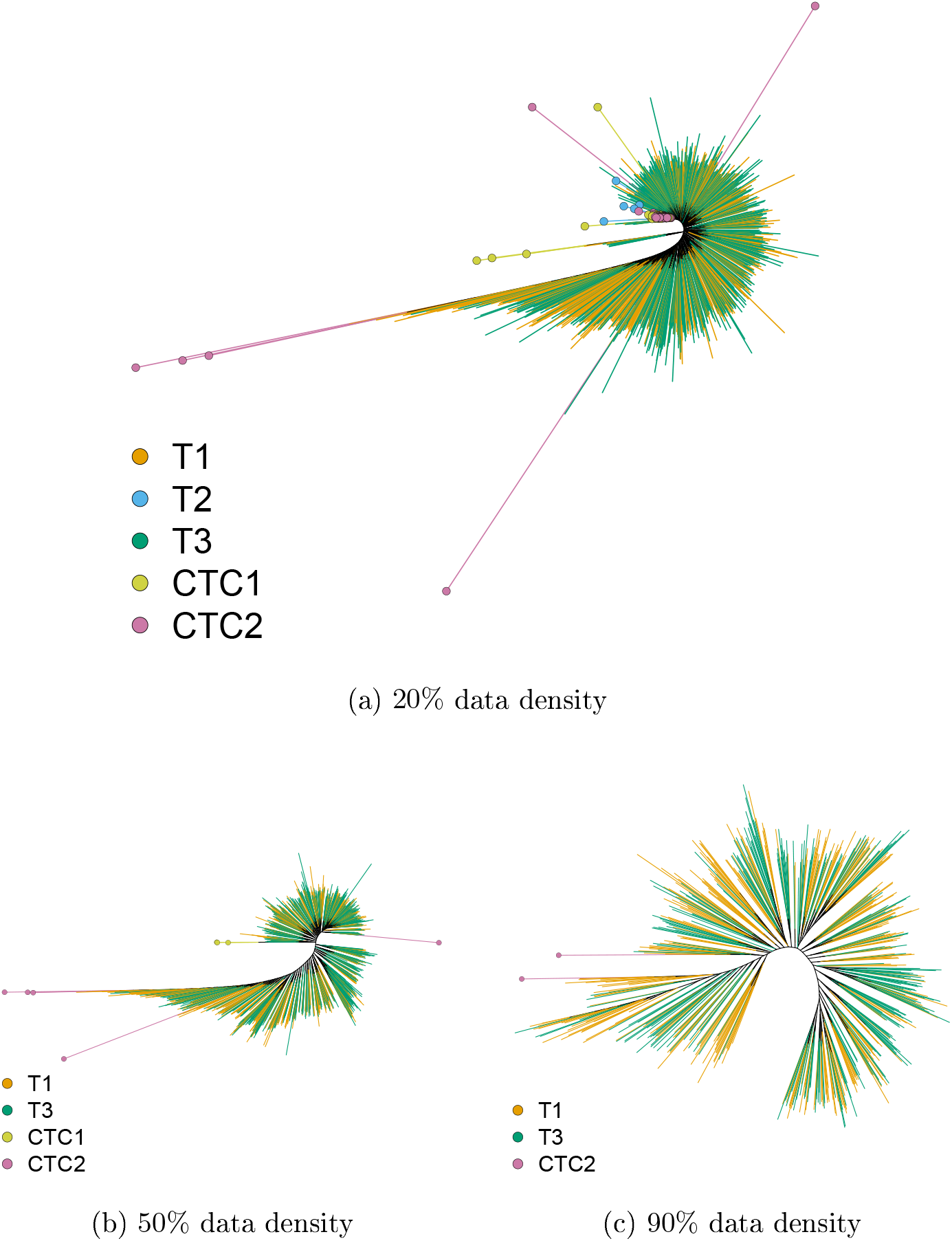
Maximum Likelihood trees from the expression data for 20% (Supplementary Figure 1a), 50% (Supplementary Figure 1b), and 90% (Supplementary Figure 1c) data density. Terminal branches are colored according to cell’s sample of origin (T1, T2, T3, CTC1, CTC2). The T2, CTC1 and CTC2 samples are marked with colored circles.

**Supplementary Figure 2.**
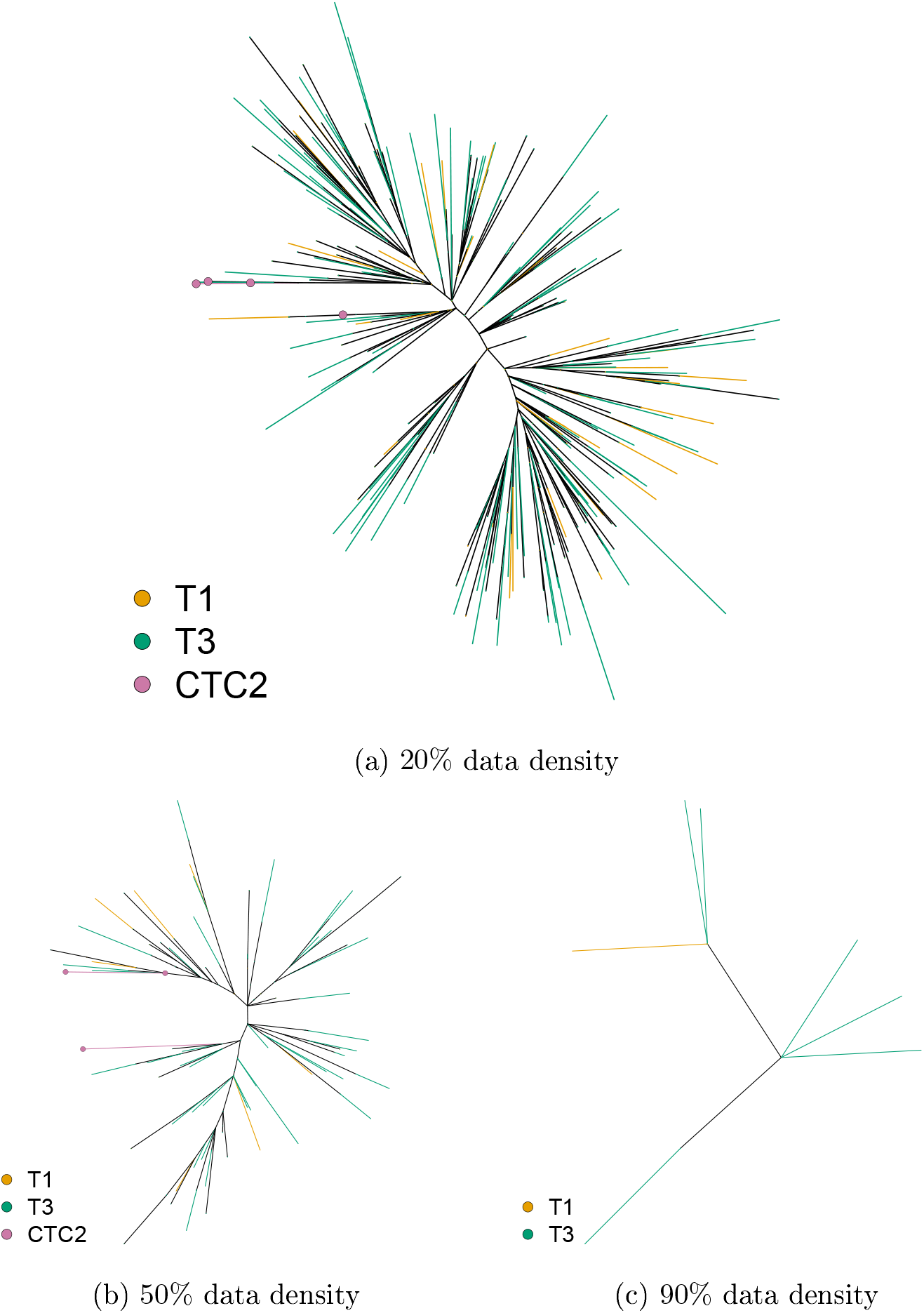
Maximum Likelihood trees from the SNV data for 20% (Supplementary Figure 2a), 50% (Supplementary Figure 2b), and 90% (Supplementary Figure 2c) data density. Terminal branches are colored according to cell’s sample of origin (T1, T2, T3, CTC1, CTC2). The T2, CTC1 and CTC2 samples are marked with colored circles.

**Supplementary Table 3.** Test of phylogenetic clustering for the datasets filtered to 20%, 50% and 90% data density. Mean Pairwise Distance (MPD) and Mean Nearest Taxon Distance (MNTD) calculated for the Maximum Likelihood (ML) and Bayesian (BI) trees from the expression and SNV data. P-values for MPD and MNTD were calculated for each sample (T1, T2, T3, CTC1, CTC2) and expected clustering for cells isolated from a single individual (T1 with CTC1, and T2 with CTC2) and to test a possible mislabeling between CTC1 and CTC2 samples (T1 with CTC2, and T2 with CTC1). Significant p-values at *α* = 0.05 after correcting for multiple comparisons using the False Discovery Rate method (Benjamini et al. 1995) are marked with an asterisk.

| Data | Groups | 20% data density |  |  | 50% data density |  |  | 90% data density |  |  |
| --- | --- | --- | --- | --- | --- | --- | --- | --- | --- | --- |
|  |  | Cells | MPD | MNTD | Cells | MPD | MNTD | Cells | MPD | MNTD |
| Expression | T1 | 701 | 1.000 | 1.000 | 688 | 1.000 | 0.996 | 329 | 0.910 | 0.089 |
|  | T2 | 11 | *0.001 | *0.001 | 0 | – | – | 0 | – | – |
|  | T3 | 806 | 0.126 | 1.000 | 758 | *0.001 | *0.001 | 262 | 0.015 | *0.002 |
|  | CTC1 | 58 | *0.001 | *0.001 | 3 | 0.156 | 0.099 | 0 | – | – |
|  | CTC2 | 51 | *0.001 | *0.001 | 5 | 1.000 | 1.000 | 2 | 1.000 | 1.000 |
|  | T1 & CTC1 | 759 | 0.992 | 0.999 | 691 | 1.000 | 0.998 | 329 | 0.910 | 0.089 |
|  | T2 & CTC2 | 62 | *0.001 | *0.001 | 5 | 1.000 | 1.000 | 2 | 1.000 | 1.000 |
|  | T1 & CTC2 | 752 | 1.000 | 1.000 | 693 | 1.000 | 1.000 | 331 | 0.972 | 0.148 |
|  | T2 & CTC1 | 69 | *0.001 | *0.001 | 3 | 0.156 | 0.099 | 0 | – | – |
| SNV | T1 | 352 | *0.001 | *0.001 | 55 | 0.066 | 0.276 | 13 | 0.221 | 0.200 |
|  | T2 | 0 | – | – | 0 | – | – | 0 | – | – |
|  | T3 | 514 | 1.000 | 0.999 | 196 | 0.795 | 0.571 | 47 | 0.801 | 0.736 |
|  | CTC1 | 0 | – | – | 0 | – | – | 0 | – | – |
|  | CTC2 | 4 | 0.520 | 0.557 | 3 | 0.892 | 0.695 | 0 | – | – |
|  | T1 & CTC1 | 352 | *0.001 | *0.001 | 55 | 0.066 | 0.276 | 13 | 0.221 | 0.200 |
|  | T2 & CTC2 | 4 | 0.520 | 0.557 | 3 | 0.892 | 0.695 | 0 | – | – |
|  | T1 & CTC2 | 356 | *0.001 | *0.001 | 58 | 0.152 | 0.558 | 13 | 0.221 | 0.200 |
|  | T2 & CTC1 | 0 | – | – | 0 | – | – | 0 | – | – |

\* significant support

### Intestinal Neuroendocrine Cancer (INC)

**Supplementary Table 4.** Test of phylogenetic clustering on the Maximum Likelihood boostrap trees reconstructed from the expression data published by Rao et al. (2020a). Mean Pairwise Distance (MPD) and Mean Nearest Taxon Distance (MNTD) calculated for the Maximum Likelihood boostrap trees reconstructed from the expression dataset containing only cancer cells and from the dataset containing all cell types. P-values were calculated for the sample of 100 bootstrap trees and 1000 posterior trees from the Maximum Likelihood and Bayesian analyses respectively, and this distribution of p-values is summarized with mean and 95% confidence interval. Significant p-values at *α* = 0.05 after correcting for multiple comparisons using the False Discovery Rate method (Benjamini et al. 1995) are marked with an asterisk.

| Groups | Cancer only |  |  | All cell types |  |  |
| --- | --- | --- | --- | --- | --- | --- |
|  | Cells | MPD | MNTD | Cells | MPD | MNTD |
| Cancer cells | 1000 | – | – | 355 | *0.001 (0.001–0.001) | *0.001 (0.001–0.001) |
| Fibroblasts | 0 | – | – | 552 | 1.000 (0.999–1.000) | 0.986 (0.967–1.000) |
| Endothelial cells | 0 | – | – | 71 | 0.797 (0.679–0.914) | 0.901 (0.811–0.977) |
| Immune cells | 0 | – | – | 22 | 0.734 (0.629–0.828) | 0.616 (0.411–0.783) |
| Metastasis | 500 | *0.001 (0.001–0.001) | *0.001 (0.001–0.001) | 500 | *0.001 (0.001–0.001) | *0.001 (0.001–0.001) |
| Primary | 500 | 1.000 (1.000–1.000) | 0.998 (0.992–1.000) | 500 | 1.000 (1.000–1.000) | 1.000 (1.000–1.000) |

\* significant support

The bootstrap analysis for the SNV dataset failed to nish after 30 days and was terminated. For this reason, the phylogenetic clustering tests for the bootstrap trees from the SNV data are omitted.

**Supplementary Table 5.**
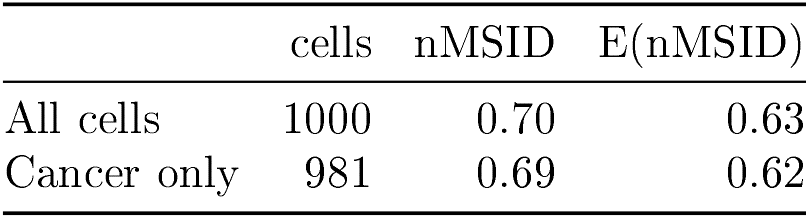
Comparison of trees reconstructed from the SNV and expression data published by Rao et al. (2020a) using nMSID. The nMSID was calculated between tree reconstructed from SNV and a tree reconstructed for expression data and compared to a expected random nMSID for trees with the same amount of taxa. For both subset (All cells and Cancer cells only), the calculated nMSID (0.70 and 0.69) were larger than expected for random trees. The information in SNV and expression values seems to be diverging.

### Gastric Cancer (GC)

**Supplementary Figure 3.**
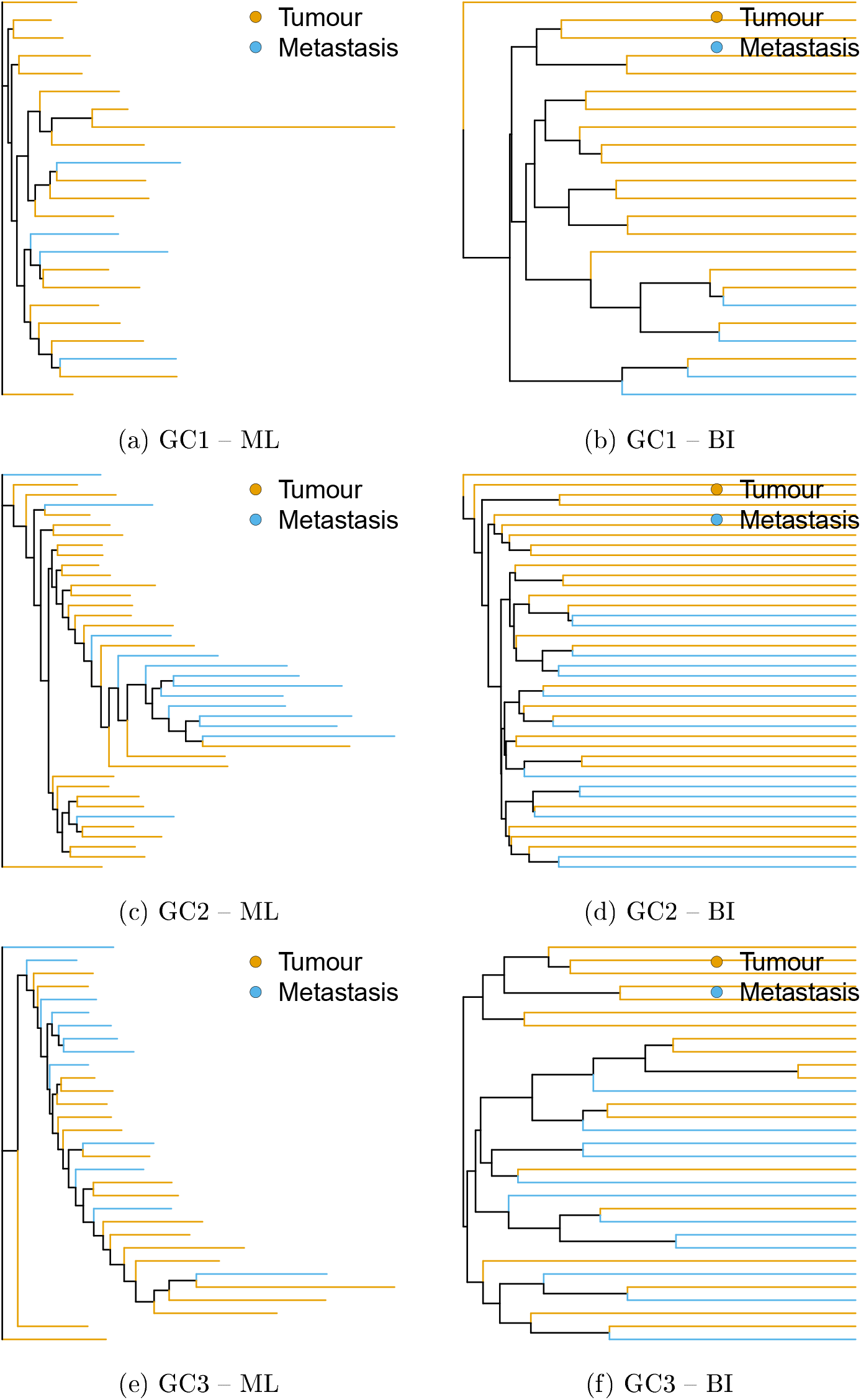
Maximum Likelihood Bayesian trees constructed from the expression data published by Wang et al. (2021). Terminal branches are colored according to cell’s sample of origin.

**Supplementary Figure 4.**
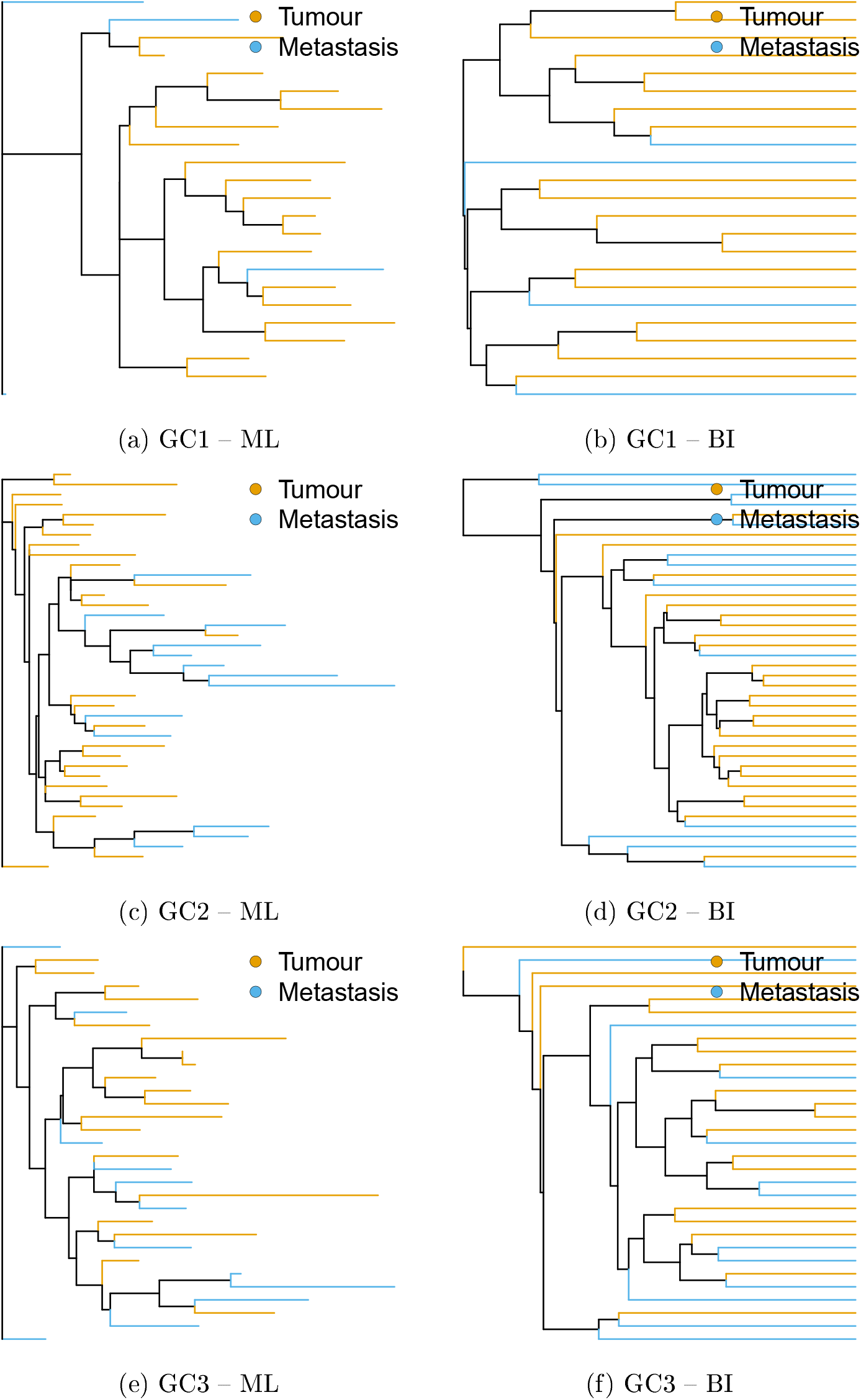
Maximum Likelihood Bayesian trees constructed from the SNV data published by Wang et al. (2021). Terminal branches are colored according to cell’s sample of origin.

**Supplementary Table 6.** Test of phylogenetic clustering on the sample of Maximum Likelihood and Bayesian trees calculated from expression and SNV data published by Wang et al. (2021). Mean Pairwise Distance (MPD) and Mean Nearest Taxon Distance (MNTD) calculated for the Maximum Likelihood boostrap trees and Bayesian posterior sample of trees reconstructed from the expression and the SNV data for patients GC1, GC2 and GC2. P-values were calculated for the sample of 100 bootstrap trees and 1000 posterior trees from the Maximum Likelihood and mean and 95% confidence interval. Significant p-values at *α* = 0.05 after correcting Bayesian analyses respectively, and this distribution of p-values is summarized with for multiple comparisons using the False Discovery Rate method (Benjamini et al. 1995) are marked with an asterisk.

| Data | Type | Groups | GC1 |  |  | GC2 |  |  | GC3 |  |  |
| --- | --- | --- | --- | --- | --- | --- | --- | --- | --- | --- | --- |
|  |  |  | Cells | MPD | MNTD | Cells | MPD | MNTD | Cells | MPD | MNTD |
| Expression | ML | Primary tumor | 19 | 0.181 (0.157–0.199) | 0.146 (0.092–0.189) | 27 | *0.001 (0.001–0.001) | *0.001 (0.001–0.001) | 19 | 0.825 (0.731–0.882) | 0.869 (0.774–0.926) |
|  |  | Lymph node | 4 | 0.816 (0.788–0.848) | 0.877 (0.723–0.979) | 13 | 0.999 (0.991–1.000) | 1.000 (0.999–1.000) | 12 | 0.143 (0.075–0.191) | 0.108 (0.041–0.187) |
|  | BI | Primary tumor | 19 | 0.661 (0.122–1.000) | 0.793 (0.402–1.000) | 27 | 0.932 (0.692–1.000) | 0.866 (0.569–1.000) | 19 | 0.394 (0.018–0.894) | 0.369 (0.002–0.814) |
|  |  | Lymph node | 4 | 0.242 (0.001–0.682) | 0.184 (0.001–0.582) | 13 | 0.033 (0.001–0.153) | 0.022 (0.001–0.101) | 12 | 0.415 (0.001–0.910) | 0.424 (0.010–0.901) |
| SNV | ML | Primary tumor | 19 | 0.198 (0.002–0.528) | 0.274 (0.005–0.682) | 27 | *0.001 (0.001–0.001) | *0.019 (0.001–0.113) | 19 | 0.594 (0.368–0.864) | 0.441 (0.074–0.900) |
|  |  | Lymph node | 4 | 0.710 (0.225–0.999) | 0.620 (0.132–0.995) | 13 | 1.000 (1.000–1.000) | 0.935 (0.598–1.000) | 12 | 0.383 (0.087–0.658) | 0.425 (0.073–0.806) |
|  | BI | Primary tumor | 19 | 0.207 (0.001–0.544) | 0.127 (0.001–0.470) | 27 | *0.001 (0.001–0.001) | 0.061 (0.001–0.191) | 19 | 0.530 (0.319–0.752) | 0.506 (0.154–0.901) |
|  |  | Lymph node | 4 | 0.762 (0.333–0.999) | 0.761 (0.342–0.999) | 13 | 1.000 (0.998–1.000) | 0.814 (0.421–0.999) | 12 | 0.470 (0.268–0.677) | 0.362 (0.031–0.701) |

\* significant support

**Supplementary Table 7.**
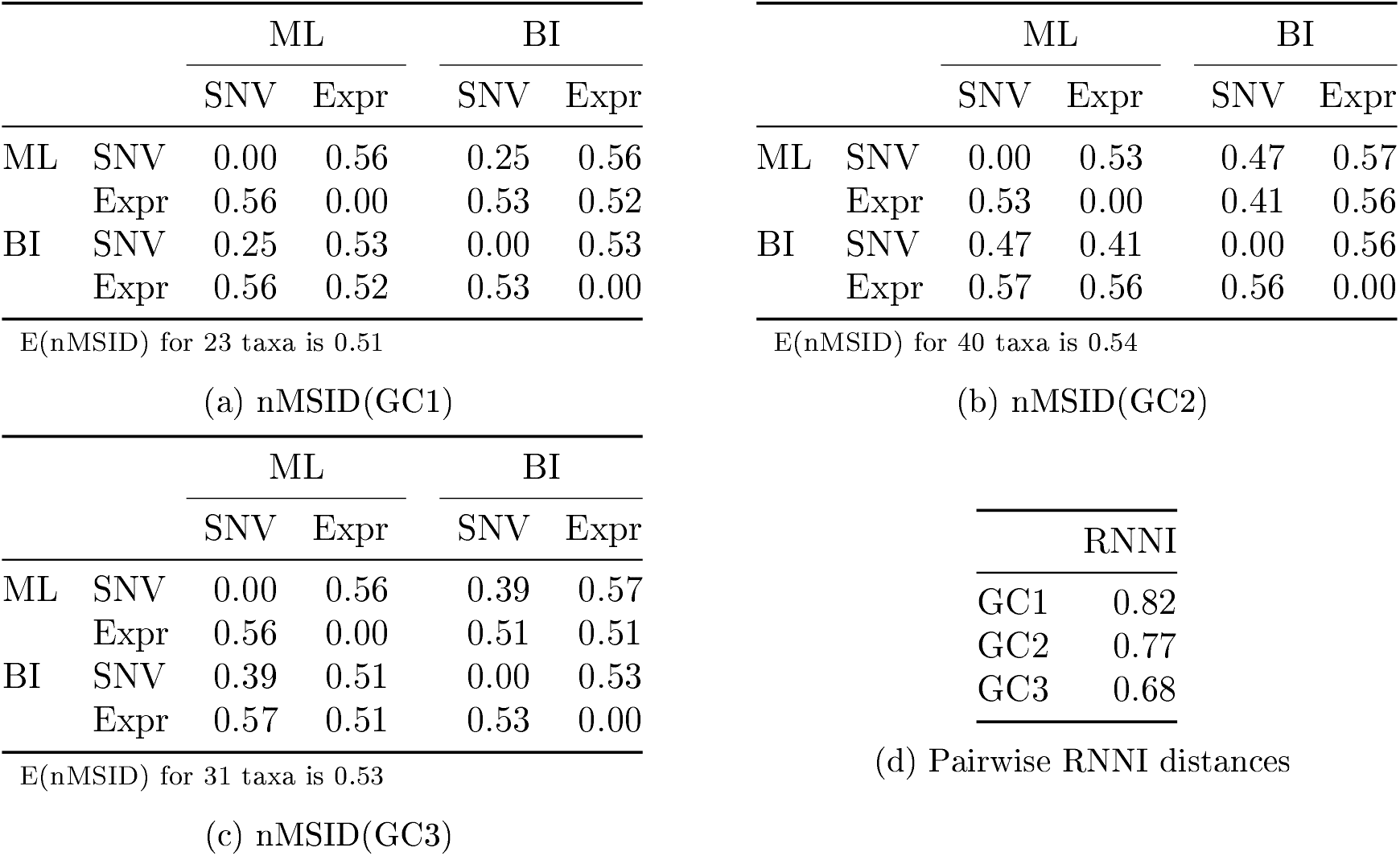
Comparison of distances between trees made from SNV and expression data from the GC dataset. Tables 7a, 7b and 7c show pairwise nMSID between the best trees from SNV and expression values from Maximum Likelihood and Bayesian Inference for patients GC1, GC2 and GC3 respectively. Table 7d shows pairwise RNNI distances between SNV and expression values for Bayesian trees for all three patients. Many of the nMSID for all three patients are close to the expected random nMSID (E(nMSID)). This suggests that there is a significant difficulty in reliably estimating a correct topology, regardless of the type of data (SNV or expression).

